# The severity of microstrokes depends on local vascular topology and baseline perfusion

**DOI:** 10.1101/2020.07.05.188565

**Authors:** Franca Schmid, Giulia Conti, Patrick Jenny, Bruno Weber

## Abstract

Cortical microinfarcts are caused by blood flow disturbances and are linked to pathologies like cerebral amyloid angiopathy and dementia. Despite their relevance for disease progression, microinfarcts often remain undetected and the smallest scale of disturbance has not yet been identified. We employ blood flow simulations in realistic microvascular networks from the mouse cortex to quantify the impact of single capillary occlusions. We reveal that the severity of a microstroke is strongly affected by the local vascular topology and the baseline flow rate in the occluded capillary. The largest changes in flow rate are observed in capillaries with two in- and two outflows. Interestingly, this specific topological configuration only occurs with a frequency of 8%, while the majority of capillaries are likely designed to efficiently supply oxygen and nutrients. Taken together, microstrokes likely induce a cascade of local disturbances in the surrounding tissue, which might accumulate and impair energy supply locally.

## 2 Introduction

As the brain’s energy storage is limited, a sustained supply of oxygen and nutrients is crucial to avoid local tissue damage. Accordingly, flow disturbances even at the level of individual vessels can result in cortical tissue lesions, so called microinfarcts [1-5]. In recent decades it has become evident that microinfarcts are linked to various pathologies, e.g. cerebral amyloid angiopathy (CAA), cognitive impairment, Alzheimer’s disease (AD) and dementia [6-10]. For example, post-mortem neuropathological studies revealed that 62% of patients with vascular dementia also suffered from cortical microinfarcts [11]. Moreover, the estimated total number of microinfarcts per individual can be as high as several thousand [12, 13].

Microinfarcts can either be ischemic (microstroke) or hemorrhagic (microhaemorrhage, cerebral microbleed) [7-9, 14]. The size of the resulting tissue lesion is comparable for microstrokes and microhaemorrhages [7]. Depending on the severity and extent of flow disturbance the diameter of documented microinfarcts ranges between 50 µm to a few millimeters [2, 4, 5, 7, 9-11]. This small size renders *in vivo* detection of microinfarcts challenging. More precisely, it has been suggested that in humans only microinfarcts >1 mm can be detected *in vivo* by high-resolution magnetic resonance imaging (MRI) or by diffusion-weighted MRI [9]. In *ex vivo* investigations the detection of microinfarcts is impeded by the *a priori* unknown location of the microinfarct. This aspect contributes to the difficulties in quantifying the microinfarct burden of an entire brain.

To overcome these challenges various animal models have been established, which allow a more refined study of the etiology of microinfarcts [7]. By occluding individual or multiple vessels via photothrombosis [1-5, 15-19] or by injecting microemboli [20-25], microstrokes can be induced and their impact on blood flow and on the surrounding neural tissue can be studied. The focus of most existing studies lies on the occlusion of penetrating vessels. This is likely due to the fact that because of their one-dimensional topology [26-28] penetrating vessels have been identified as the “bottleneck of perfusion” [1]. Consequently, the occlusion of a single penetrating vessel causes significant tissue damage [1-5, 16] as well as functional deficits and cognitive impairment [2, 5]. Additionally, the appearance of these microinfarcts is comparable to microinfarcts in humans [2] and it has been suggested that human microinfarcts are often centered around penetrating arterioles [29, 30]. As such, the occlusion of individual penetrating arterioles has been proven to be a suitable animal model to study the impact and progression of microinfarcts.

Less attention has been given to occlusions of descending arteriole (DA) offshoots and capillaries. While the occlusion of DA offshoots causes a maximal infarct volume of 0.8 nl (275 times smaller than for DA occlusions), no tissue damage could be detected for the occlusion of capillaries >2 branches apart from the DA [2]. However, the effect of anesthesia on these results remains unknown. This is because anesthesia can act as a vasodilator and tends to increase red blood cell (RBC) flux and tissue oxygenation [31, 32]. Importantly, it has also been shown that single capillary occlusion causes flow reversals and RBC speed reductions of up to 90% in the vessels downstream of the occluded capillary [16].

Additionally, single capillary occlusion can induce the formation and alter the morphology of amyloid-beta (Aβ) plaques [17], which are related to AD and CAA. Interestingly, Aβ deposits are frequently observed at the core of larger microinfarcts [33, 34] and have been associated with blood flow disturbances, which ultimately might lead to microhaemorrhages [6]. Moreover, a higher Aβ prevalence has been shown to negatively affect tissue clearance via the perivascular system [35-37] and therewith it might further aggravate the build-up of Aβ deposits. Taken together, even if a single capillary occlusion might not directly lead to local tissue damage, it disturbs local blood flow and tissue clearance and thus might be an important factor in the development and progression of larger disturbances and pathologies.

Further studies have investigated the effect of simultaneously occluding multiple capillaries or multiple microvessels of larger caliber [15, 25]. Underly et al. [15] showed that the occlusion of ∼10 proximal capillaries via photothrombosis leads to deterioration of the blood brain barrier (BBB). Additionally, in response to the injection of ∼1500 polystyrene microspheres (10 or 15 µm) via the carotid artery Lam et al. [25] temporarily observed small hypoxic areas and local synaptic pruning. The accumulation of occluded capillaries can also influence the total perfusion of the cortical vasculature. In an AD mouse model, Cruz Hernandez et al. [38] showed that with 2% of capillaries stalled, cortical cerebral blood flow is reduced by ∼5%.

These aspects underline the crucial need for an in-depth quantification of blood flow changes in response to single capillary occlusion to better understand the role of these small flow disturbances for the development of the aforementioned pathological conditions. Moreover, it is important to identify the smallest scale of disturbance that causes a response in the closely connected vessels and the proximal neural tissue. Quantifying the smallest scale of disturbance is a prerequisite for the correct interpretation of disturbances and changes observed on a larger scale. It is also important to note that by looking at the smallest possible scale of occlusion valuable insights on the robustness of perfusion can be gained and our knowledge of topological characteristics of cortical capillary beds can be extended.

Here, we employ blood flow simulations in realistic microvascular networks from the mouse cortex [28, 39, 40] to study the impact of single capillary occlusions on the perfusion of the cortical microvasculature. Using an *in silico* approach comes with several advantages. First of all, it is challenging to monitor blood flow changes *in vivo* with single vessel resolution in multiple vessels or even entire vascular networks simultaneously. This problem is even more pronounced, if the focus is on blood flow changes in the capillary bed, which is highly interconnected [26, 28, 39, 41-43] and in which the flow field is highly heterogeneous and fluctuating [39, 40, 44-47]. For a given numerical modeling setup the flow field is known in every vessel at any given moment. Secondly, *in silico* studies allow us to investigate the impact of single capillary occlusions in an isolated manner. This is in contrast to *in vivo* analyses, where a capillary occlusion will always be accompanied by a response from directly neighboring cells (e.g. endothelial cells, mural cells, microglia).

By studying flow changes in response to the occlusion of 96 different capillaries, we reveal that the severity of a microstroke strongly depends on the local vascular topology and the baseline flow rate in the occluded capillary. More precisely, in the worst-case scenario the flow rate drops by as much as 80% in the direct vicinity of the occluded capillary. As well as this, a microstroke locally reduces the number of available flow paths between DAs and ascending venules (AVs). This aspect might also play an important role for the up-regulation of blood flow during neural activation, since the ability of the microvasculature to adapt to local changes in energy demand might be impaired in the disturbed flow field. Our results further indicate that the different local vascular topologies are not only relevant for the severity of the microstroke, but that the different topologies might in fact fulfill distinct functional tasks. We postulate that there is a topological difference between capillaries responsible for the distribution of blood and capillaries responsible for supplying oxygen and nutrients to the cortical tissue.

In summary our work provides an in-depth quantification of flow changes in response to single capillary occlusions and reveals novel topological characteristics of the cortical micro-vasculature. Our results give valuable insights into the role of microinfarcts, which are relevant for future *in vivo* studies as well as for the progression of various pathologies in general.

## 3 Results

The following results are based on time-averaged blood flow simulations in realistic microvascular networks (MVNs). The realistic MVNs and the simulation framework have been introduced in previous publications [28, 39, 48]. A brief description of both is provided in the Methods. In total we performed 96 microstroke simulations to investigate the impact of a microstroke on local perfusion and to reveal key factors for the severity of the microstroke. To induce a microstroke we constrict the diameter of the microstroke capillary (MSC) to 0.01 µm, which reduces the flow rate in the MSC to < 10 ^−10^ *μm*^3^*ms*^−10^. Details regarding the selection of MSCs and the computation of the relative flow changes Δ*q*_*ij*_ are provided in the Methods and in Supplementary Table 1.

### 3.1 The severity of a microstroke is governed by the local vascular topology

Microvascular bifurcations are either divergent or convergent. Thus, depending on the bifurcation types at the source and the target vertex of the MSC, four topological configurations are possible at the MSC (Figure 1a-d). To investigate the impact of the topological configuration on the severity of the microstroke we perform eight microstroke simulations per configuration. Based on the time-averaged flow field before and after stroke we compute the thresholded relative flow change Δ*q*_*ij*_ for each vessel (Methods).

**Figure 1.**
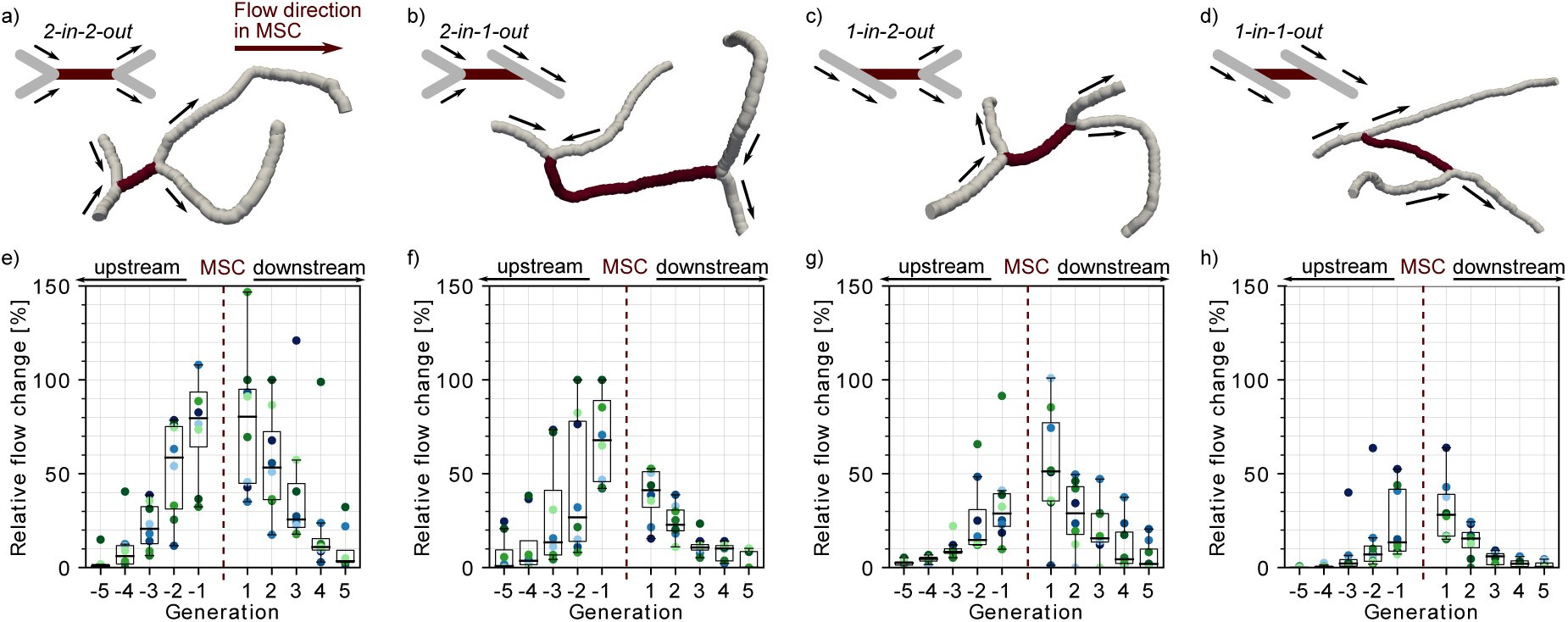
Impact of the local vascular topology on the severity of a microstroke. a)-d) Illustration of the four possible topological configurations at a microstroke capillary (MSC). For each topological configuration a schematic (upper left) and a realistic example (lower right) is provided. The MSC (dark) and its adjacent vessels (grey, generation −1 and 1) are depicted. The arrows show the flow direction. e)-f) Average relative change in flow rate for ***Δq***_***ij***_ capillaries up- and downstream of the MSC. For each topological configuration the flow field for eight microstrokes has been computed. The average relative change per generation for each of the eight simulations is depicted by the color-coded spheres. Note that the number of up- and downstream vessels per generation varies between MSCs. The boxplots are based on the data for each generation.

In a first step, we analyze the relative flow changes Δ*q*_*ij*_ in the vessels in up to five generations up- and downstream of the MSC. Figure 1e-g shows that the relative changes are larger for MSCs fed by two upstream vessels and for MSCs feeding two downstream vessels. For the *worst-case scenario*, i.e. MSCs with a convergent bifurcation upstream and a divergent bifurcation downstream (*2-in-2-out*, Figure 1a and e), the median relative change is still >20% at generation ±3 from the site of occlusion. In contrast for the *best-case scenario*, i.e. MSCs with a divergent bifurcation upstream and convergent bifurcation downstream *(1-in-1-out*, Figure 1d and h), the median relative change is below 10% at generation ±3. The differences between *2-in-2-out* and *1-in-1-out* are even more pronounced for vessels of generation ±1. Here, the median relative drop in blood flow is as large as 80% for *2-in-2-out*, while it is only 15% and 40% for generation −1 and 1 respectively for *1-in-1-out*.

*2-in-1-out* and *1-in-2-out* are intermediate MSC-types (Figure 1 b-c and f-g). For example, on the upstream side *2-in-1-out* experiences relative changes comparable to *2-in-2-out*, while on the downstream side the trends correspond to the ones observed for *1-in-1-out*. It is noteworthy that for the given example the changes on the upstream side are smaller than for *2-in-2-out* and larger on the downstream side in comparison to *1-in-1-out*.

Analyzing the flow direction changes reveals that the occlusion of a *2-in-2-out* mostly leads to a flow reversal in one of the two vessels at generation −1 and 1 (Supplementary Figure 1a). This is plausible, because from a fluid dynamical point of view there are only two possible outcomes for a *2-in-2-out* occlusion. The first is the observed flow reversal at generation ±1. The second would be the complete cessation of flow in generation ± 1, which would lead to a significantly larger infarct volume and thus would increase the severity of the microstroke. The cessation of flow in generation ±1 has only been observed once downstream of a *2-in-2-out* and twice upstream of a *2-in-1-out*. In all three scenarios a very specific topology was identified at generation 1 (*2-in-2-out*)/−1 (*2-in-1-out*), where the two generation ±1 vessels are connected to the same generation ±2 vessel (Supplementary Figure 1b). We conclude that the cessation of flow in vessels adjacent to the MSC is rare and that the less severe flow reversal is more common. Interestingly, the occlusion of a *1-in-1-out* capillary does not cause any flow direction changes.

To predict oxygen and nutrient supply in the tissue around the MSC it is important to account for changes in capillaries, which are in the direct vicinity of the MSC but not directly upstream or downstream of the MSC. Therefore, we define an analysis box around the MSC and compute its total inflow before and after stroke (Figure 2a-d, Methods). The initial analysis box volume is set to 200,000 µm^3^ and is chosen such that each MSC fits into the initial box volume and that the box has at least five inflows. The box volume is increased progressively and the relative inflow difference is recomputed. This analysis allows us to comment on the reduction in perfusion of a tissue volume around the MSC capillary. Moreover, it provides an estimate of the tissue volume, which is affected by a reduced perfusion in response to the microstroke.

**Figure 2.**
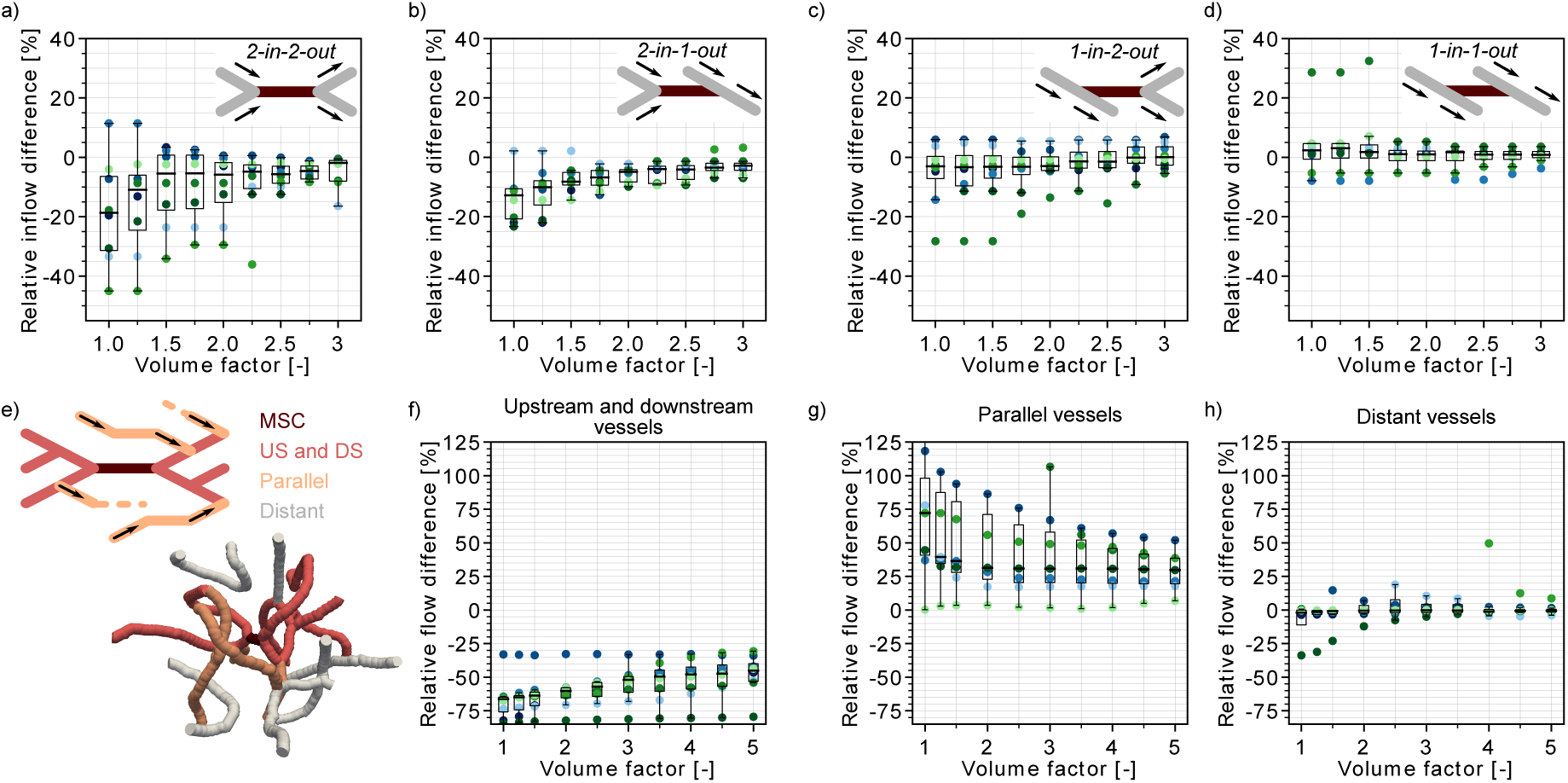
Flow reduction in an analysis box around the microstroke capillary (MSC). a)-d) Relative inflow difference for an increasing box volume around the MSC for the four MSC-types. a) *2-in-2-out*, b) *2-in-1-out*, c) *1-in-2-out* and d) *1-in1-out*. The initial box volume, i.e. volume factor = 1, is 200,000 µm^3^. The relative inflow difference is computed by adding up the inflows across the borders of the box for the baseline and the stroke simulation (Methods). e) Upper panel: Schematic to introduce the concept of vessels parallel to the MSC (Methods). Lower panel: Realistic example of the edges in a box volume of 1,000,000 µm^3^, i.e. volume factor = 5. US: upstream. DS: downstream. f)-h) Relative total flow difference for an increasing box volume around a *2-in-2-out* MSC for upstream and downstream vessels (f), parallel vessels (g) and distant vessels (h). The relative total flow difference is calculated by adding up the length-weighted flow for the baseline and the stroke simulation (Methods). The eight microstrokes per topological configuration are depicted by the color-coded spheres. The boxplots are based on the data for each generation.

In line with the relative flow changes in the upstream and downstream vessels (Figure 1e-h) we observe the largest inflow reduction for *2-in-2-out* (Figure 2a). For the initial box volume, i.e. a volume factor of 1.0, the median inflow reduction is as large as −19%. For a volume factor of 1.5 the median inflow reduction already drops to −5.5%. However, it is not until a volume of 600,000 µm^3^ is reached (volume factor: 3.0) that the median inflow reduction approaches 0%. For MSC-type *2-in-1-out* the median inflow reduction for a volume factor of 1.0 is −12% (Figure 2b). The tissue around MSC-types *1-in-2-out* and *1-in-1-out* does not experience significant changes in total inflow. Here, for all volume factors the median inflow difference is smaller than 5%.

Importantly, the resulting inflow reduction in the analysis box is also affected by the topological connectivity around the MSC and the redistribution of flow in response to a microstroke. These aspects become apparent if we compare the relative flow changes in vessels with different topological positions with respect to the MSC. We discern three topological positions: 1) vessels that are directly upstream and downstream of the MSC, 2) vessels that run parallel to the MSC and 3) vessels that do not belong to the first two categories, i.e. distant vessels (Figure 2e, Supplementary Figure 2e, Methods).

Supplementary Figure 2a shows that for up to a volume factor of 2.5, more than 50% of the vessels in the analysis box are directly upstream or downstream of the MSC. In these vessels we have a significant flow reduction (Figure 2f). In contrast, in the vessels that run parallel to the MSC we observe an increase in flow (Figure 2g). This clearly indicates that during a microstroke the flow is redistributed to pathways parallel to the MSC. However, only approximately ∼20% of capillaries are parallel in the analysis box (Supplementary Figure 2b) and consequently we still observe an overall flow reduction in the analysis box. In the third vessel category, the distant vessels, the median relative flow difference is smaller than 2% for all volume factors (Figure 2h). This result confirms that the impact of a microstroke is most pronounced in vessels that are directly connected to the MSC.

Worth of a not is that for a tissue volume of 600,000 µm^3^ (i.e. volume factor = 3) at least 40% of vessels are distant and parallel in the analysis box (Supplementary Figure 2d). This topological configuration might be beneficial for the robustness of perfusion of the tissue volume around the MSC. Because even if the total inflow decreases in the tissue volume around the MSC, there is always a fraction of vessels in the analysis box that are not significantly affected by the microstroke (distant vessels) and vessels which experience an increase in flow in response to the microstroke (parallel vessels). Therewith, an even larger drop in overall perfusion can be avoided and a minimum remaining perfusion can be ensured.

### 3.2 The baseline MSC flow rate increases the area of impact of a microstroke

Our results demonstrate that the local vascular topology plays a crucial role in the severity of a microstroke. To identify further structural and functional characteristics relevant to the level of flow change in response to microstroke, we repeat our analysis for eight additional vessel subsets. We look at the impact of the baseline flow rate in the MSC (case 5), the cortical depth (case 8-12) and the distance to the penetrating vessels (case 6-7). An overview of the selection criteria for each vessel subset is provided in Supplementary Table 1. For each case eight microstroke simulations have been performed. In this study we focus on *2-in-2-out* MSCs because we expect the largest changes here.

Of all additionally tested cases only the baseline flow rate in the MSC affects the severity of the microstroke. Although the relative changes up to generation ±3 do not differ significantly for a low and a high baseline flow rate, the relative change at generation ±5 is still ∼10% for the MSC with the high baseline flow rate (Supplementary Figure 3a-b). This result is confirmed by the analysis of the total inflow change into the analysis box around the MSC (Supplementary Figure 3c-d). With a median inflow reduction of −37% for the initial box volume, the inflow reduction is nearly twice as large for the high baseline flow capillaries. Moreover, the median inflow reduction is still as large as −14% for a box volume of 600,000 µm^3^, i.e. a volume factor of 3.

Only very small differences can be observed for the occlusion of MSCs at different cortical depths (Supplementary Figure 4a-d). These differences can probably be explained by a general decrease in flow rate over depth and a more homogeneous flow field for deeper cortical layers [39, 46, 49, 50]. Moreover, our results suggest that the position of the MSC along the path between DA and AV does not play a significant role in the severity of the microstroke (Supplementary Figure 5).

### 3.3 MSC-type *1-in-1-out* supplies the largest tissue volume and is the most frequent MSC-type

The significant impact of the different topological configurations on the severity of the microstroke raises questions about the frequency of occurrence and the distribution of the different MSC-types in a realistic MVN. Note that the following investigations are based on two realistic MVNs, which jointly encompass a tissue volume of ∼3.6 mm^3^ and which contain 31,400 vessels.

Interestingly, the *worst case scenario*, i.e. MSC-type *2-in-2-out*, only occurs with a frequency of 8%, while the *best-case scenario*, i.e. MSC-type *1-in-1-out*, represents 43% of all possible MSCs (Figure 3b). Moreover, the median supplied tissue volume of *1-in-1-out* is 79% larger than the supplied tissue volume of *2-in-2-out* (Figure 3c, Methods). Consequently, a total of 53% of the tissue is supplied by *1-in-1-out* capillaries with only 6% supplied by *2-in-2-out* capillaries. This also becomes apparent in Figure 3f where the tissue volume of realistic MVN1 is color-coded based on the MSC-type by which it is supplied. The differences in the median supplied tissue volume are partly caused by the larger median vessel length of *1-in-1-out* capillaries (Supplementary Figure 6b).

**Figure 3.**
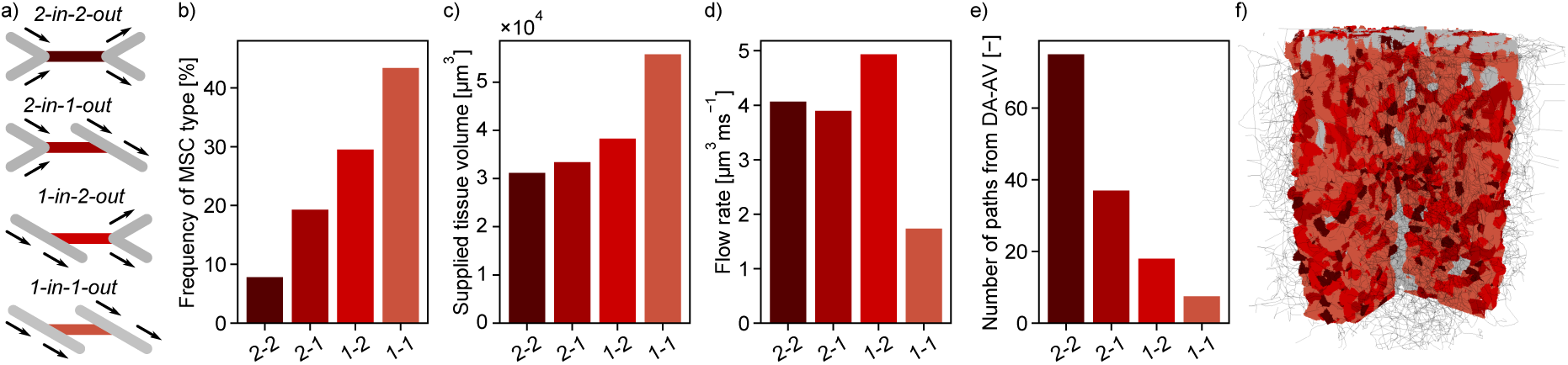
Characteristics of the four topological configurations at a microstroke capillary (MSC). a) Schematic of the four topological configurations at a MSC. The MSC is color coded in accordance with subfigures b)-f). b) Frequency of occurrence of the four MSC-types. c) Median supplied tissue volume for the four MSC-types (Methods). d) Median flow rate for the four MSC-types. e) Median number of unique paths leading through a MSC from the descending arteriole (DA) to the ascending venule (AV). f) Grid representation of the tissue in which the realistic microvascular network (MVN) is embedded. The tissue points are color-coded based on the closest MSC-type. Abbreviations of the four MSC-types: 2-2: *2-in-2-out*, 2-1: *2-in-1-out*, 1-2: *1-in-2-out*, 1-1: *1-in-1-out*. The statistics are based on all capillaries that fulfill the general selection criteria described in the Methods. The fifth selection criterion is less strict for the current analysis, i.e. the capillary only has to be one segment apart from the DA/AV, and the sixth criterion is not applied. This results in 4818 and 8544 capillaries for analysis for MVN1 and MVN2, respectively. First the median within each MVN is computed and subsequently the average across the two MVNs.

The small number of *2-in-2-out* capillaries and the small flow reduction for the frequent MSC-type *1-in-1-out* suggest that the cortical microvasculature is inherently robust to the occlusion of a single capillary. The significant differences in the supplied tissue volume further underline this aspect.

Figure 3d shows that the median flow rate in a *2-in-2-out* capillary is 2.3 times larger than in a *1-in-1-out*. As a higher baseline flow rate increases the area of impact of a microstroke, we conclude that these differences further contribute to the severity of a microstroke in a *2-in-2-out* configuration. We hypothesize that different local topological configurations might fulfill different tasks in microvascular blood supply. While *2-in-2-out* capillaries might be more relevant for distributing blood in the cortical microvasculature, *1-in-1-out* capillaries are likely designed to robustly deliver oxygen and nutrients to the cortical tissue. This hypothesis is strengthened by the number of unique paths going from DA to AV through the different MSC-types (Figure 3e, Methods). While for *1-in-1-out* we only have 8 unique paths connecting DA and AV, for *2-in-2-out*, we have 75 unique paths. The observed trends are consistent between the two MVNs (Supplementary Figure 7). However, it is noteworthy that the overall average flow rate is larger in MVN2 and the number of paths per vessel is significantly larger in MVN1. The latter is likely caused by a higher density of penetrating vessels in MVN2, which reduces the number of highly interconnected flow paths through the capillary bed.

Subsequently, we asked whether the frequency of MSC-types varied over cortical depth (Supplementary Table 2) or along the pathway between DA and AV (Supplementary Table 3). The latter investigation was performed by varying the constraint on the minimum distance between MSC and DA and AV, respectively. For both investigations no significant differences in the frequency of the MSC-type could be detected. As previously mentioned, the median flow rate decreases significantly over cortical depth (−66% and –80% for MVN1 and MVN2, respectively). This is consistent for all MSC-types (Supplementary Figure 8f-j). The maximal difference in the median supplied tissue volume between analysis layers (ALs) is 36%. However, no consistent trend as to how the supplied tissue volume changes over depth can be identified (Supplementary Figure 8k-o). Important to note, is that the largest supplied tissue volume is found for *1-in-1-out* capillaries in all ALs and the relative supplied tissue volume (Supplementary Figure 8p-t) does not vary significantly with depth. Consequently, our conclusion holds that *1-in-1-out* capillaries might be key capillaries for nutrient and oxygen discharge.

To further elucidate the distribution of the four MSC-types along the capillary pathway we compute the minimum and median distance for each capillary to a DA and AV main branch (Supplementary Figure 6). Keep in mind that for nearly all capillaries there are multiple paths leading to the DA and AV. Here, we analyze the minimum and the median distance of the set of all available paths. Both the minimum and the median distance show that *1-in-2-out* tends to be the MSC-type closest to the DA, while *2-in-1-out* is closest to the AV. From a topological point of view this is plausible because blood is distributed to the capillary bed (*1-in-2-out*) close to the DA and it is recollected at the venule end of the capillary bed (*2-in-1-out*). Notably, 76.5% of all paths between DA and AV contain each MSC-type at least once (MVN1: 92%, MVN2: 61%). The MSC-type, *2-in-2-out*, is that which is most frequently missing along a path between DA and AV.

### 3.4 A microstroke locally reduces the number of available flow paths

To further investigate the redistribution of flow during a microstroke we analyze the number of flow paths leading from DA to AV. To this end, we follow the flow downstream from the main branch of a DA until it reaches an AV main branch (Figure 4b, Methods). Importantly, due to the finite size of the MVN, various flow paths do not start at a DA or do not end at an AV. These flow paths are not considered (Methods).

**Figure 4.**
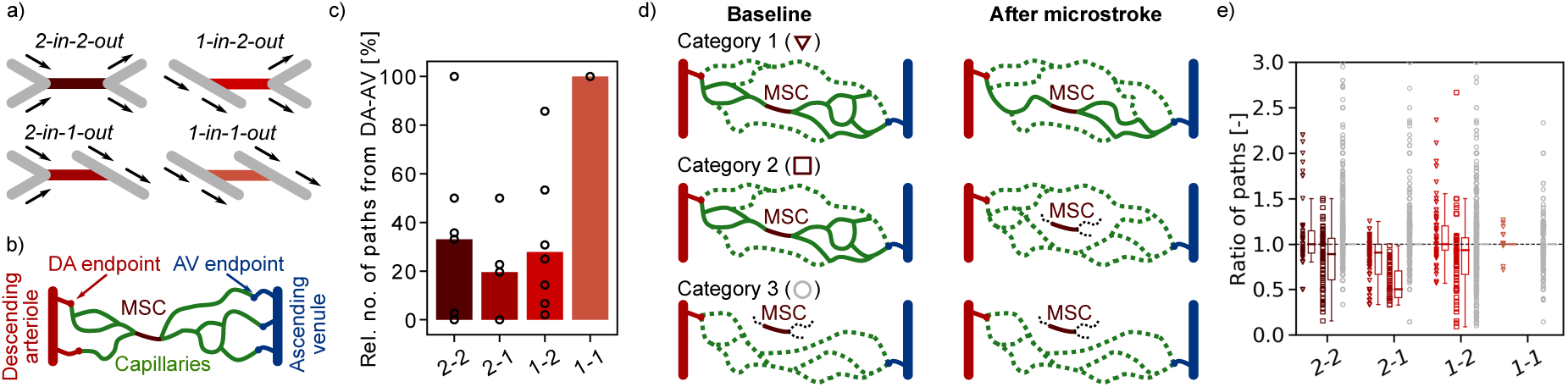
Changes in flow paths in response to a microstroke in MVN1. a) Schematic of the four topological configurations at the microstroke capillary (MSC). The MSC is color coded in accordance with subfigures c) and e). b) Schematic to introduce *DA-AV-endpoint-pairs* and the concept of flow paths through a MSC. The illustrated example has 15 flow paths going through the MSC and 6 *DA-AV-endpoint-pairs*. Only paths through the MSC are depicted. c) Relative number (Rel. no.) of flow paths through the MSC, which connect a descending arteriole (DA) to an ascending venule (AV). The relative number is computed with respect to the baseline case (Methods). The bar plot depicts the median for each MSC-configuration. The spheres show the relative number of flow paths for the eight microstrokes for each MSC-type. d) Schematic to introduce the three categories of *DA-AV-endpoint-pairs* (Methods). Each of the subplots shows all flow paths between one *DA-AV-endpoint-pair*. Flow paths that do not go through the MSC are labeled by the dotted line. e) Ratio of the number of flow paths between *DA-AV-endpoint-pairs* of different categories (see d) and for different MSC-types. Each marker represents the ratio for a unique *DA-AV-endpoint-pair*. The ratio is obtained by dividing the number of flow paths between a *DA-AV-endpoint-pair* pre stroke by the number of paths post stroke (Methods). The raw data for each category of all eight microstroke cases per MSC-type is shown by the scatter points and summarized in the boxplot to the right of it. Triangles: *DA-AV-endpoint-pairs* of category 1, i.e. at least one path through the MSC before and after stroke. Squares: *DA-AV-endpoint-pairs* of category 2, i.e. only before stroke is the *DA-AV-endpoint-pair* connected by a path through the MSC. Circles: *DA-AV-endpoint-pairs* of category 3, i.e. none of the paths connecting the DA- and AV-endpoint go through the MSC. 0.08% of the data are not displayed (ratio > 3). Abbreviations of the four MSC-types: 2-2: *2-in-2-out*, 2-1: *2-in-1-out*, 1-2: *1-in-2-out*, 1-1: *1-in-1-out*.

To study the role of individual capillaries for the distribution of flow we count the number of flow paths going through a predefined capillary (Figure 4b). As expected the total number of flow paths going through the MSC drops significantly during stroke for all MSC-types except for type *1-in-1-out* (Figure 4c). Note that for most cases some flow paths through the MSC remain. However, as previously mentioned, the flow rate in the MSC drops below 10^−10^*μm*^3^*ms*^−10^ after stroke and consequently, the remaining paths carry a negligible amount of flow. As there is no significant drop in the total number of unique flow paths through the MVN we conclude that a microstroke mostly affects the closely connected vessels of the MSC but not the overall flow field of the MVN (Supplementary Figure 9b).

Next, we analyzed the change in the number of unique flow paths going through: 1) capillaries upstream and downstream of the MSC (up to generation 3), 2) parallel to the MSC and 3) distant to the MSC (Figure 2e, Methods). Based on the results presented in Figure 2f-h, where we detected an increased flow in the parallel vessels, we expected to see an increase in the number of paths going through the parallel vessels. However, no consistent trend could be observed for the three different vessel categories (Supplementary Figure 9c). This suggests that an increase in flow does not necessarily cause an increase in the number of flow paths through the respective capillary.

To further extend our understanding of flow redistribution in response to a microstroke we will now examine the number of possible flow paths between a given DA- and AV-endpoint before and after stroke. In a first step, we compare the total number of possible DA-AV-endpoint-combinations (*DA-AV-endpoint-pairs*). Although, we observe some differences in the number of unique *DA-AV-endpoint-pairs* (Supplementary Figure 9d), the overall change is small (< 1.1%) with respect to the total number of possible *DA-AV-endpoint-pairs* (2,380 *DA-AV-endpoint-pairs* at baseline).

To study the flow paths between given *DA-AV-endpoint-pair*s we introduce three categories to classify *DA-AV-endpoint-pair*s (Figure 4d): 1) before and after stroke there is at least one path that leads from the DA- to the AV-endpoint through the MSC, 2) only before stroke is there at least one path that leads from the DA- to the AV-endpoint through the MSC and 3) none of the paths between the given *DA-AV-endpoint-pair* go through the MSC. It is only for the MSC-type *2-in-1-out* that we observe a decrease in flow paths between *DA-AV-endpoint-pairs* of category 1 (Figure 4e). For MSC-types *2-in-2-out* and *1-in-2-out*, increases as well as decreases in the number of flow paths occur and consequently, the median of the ratio of flow paths is close to 1. However, it is important to note that only a small fraction of *DA-AV-endpoint-pairs* belong to category 1 (Supplementary Figure 9e). To be more precise, only ∼1/3 of the *DA-AV-endpoint-pairs* that are connected by a path through the MSC during baseline also have a path through the MSC during stroke. The remaining 2/3 lose their path through the MSC and consequently belong to category 2 *DA-AV-endpoint-pair*s.

For *DA-AV-endpoint-pairs* of category 2 the trends between the MSC-types are more consistent. Here, we clearly note a decrease in the number of available flow paths between the respective *DA-AV-endpoint-pairs*. The decrease in the number of pathways between *DA-AV-endpoint-pairs* of category 2 shows that a microstroke locally reduces the number of available paths and that the flow is not inherently redistributed in a way that preserves the number of flow paths. No trend was observed for the changes in *DA-AV-endpoint-pairs* of category 3 and for all *DA-AV-endpoint-pairs* of MSC-type *1-in-1-out*.

Taken together, our results suggest that a microstroke locally causes a drop in the number of available flow paths between DA and AV. However, as shown in the preceding sections, the flow rate likely increases along some of the remaining paths.

### 3.5 The minimum distance between an arteriole-sided and a venule-sided capillary point is on average 58 µm

It is well established that the oxygen partial pressure in capillaries shortly downstream of DAs is higher than in capillaries just upstream of AVs [50, 51]. Moreover, it has been suggested that the tissue supplied by venule-sided capillaries might be more susceptible to hypoxia in the case of blood flow disturbances or during neural activation [51-53]. Consequently, the arrangement of arteriole-sided and venule-sided capillaries with respect to each other might be an important topological feature for the robustness of oxygen and nutrient supply. A convenient way to avoid local hypoxia could be obtained by a topological structure where arteriole-sided and venule-sided capillaries are positioned in close proximity to each other.

To investigate the arrangement of arteriole-sided and venule-sided capillaries with respect to each other we introduce the *AV-factor*. The *AV-factor* for each capillary is computed by identifying all paths leading from the capillary to all possible DA-endpoints and to all possible AV-endpoints. The *AV-factor* is subsequently calculated from the median distance to all DA/AV-endpoints (Methods). The *AV-factor* is close to 0 if the capillary is close to a DA and is close to 1 if the capillary is adjacent to an AV. We define arteriole-sided capillaries as capillaries with an *AV-factor* < 0.5 and venule-sided capillaries with an *AV-factor* >= 0.5. The following investigations have been performed in MVN1 and MVN2.

In an initial analysis we compute the shortest distance between a point along a venule-sided capillary to a point along an arteriole-sided capillary. Of interest to note, is that the median shortest distance to an arteriole-sided capillary is 58 µm. Therewith, the shortest distance between a venule-sided capillary point and an arteriole-sided capillary is only ∼8 µm larger than the average distance between two capillaries [51, 54]. Subsequently, we analyzed the average *AV-factor* surrounding a venule-sided capillary point. For 44% of all venule-sided capillary points, the average *AV-factor* in a sphere of 50 µm radius was smaller than the *AV-factor* at its center point. This implies that these points have multiple arteriole-sided points nearby, which potentially act as backup for oxygen and nutrient supply. The relative difference between the *AV-factor* of the center point and the mean of all points in the 50 µm sphere was −4.1%.

Taken together, the shortest distance of 58 µm to an arterial-sided capillary and the average decrease of the average *AV-factor* in a 50 µm sphere around a venule-sided capillary point suggest that arteriole-sided capillaries are well distributed throughout the network. Nonetheless, further studies are necessary to estimate whether proximal arterial-sided capillaries help to avoid hypoxic tissue areas in the vicinity of venule-sided capillaries during flow disturbances. Moreover, it has to be kept in mind that in the rodent cortical vasculature, AVs outnumber DAs [26]. This is in contrast to the primate vasculature where DAs are approximately twice as frequent as AVs [26, 46, 55, 56]. As such, the current conclusion needs to be verified for different species.

## 4 Discussion

By performing blood flow simulations in realistic MVNs for a large number of single capillary occlusions we show that the severity of a microstroke depends on the local vascular topology and on the baseline flow rate in the occluded capillary. The largest impact is observed if capillaries with two in- and two outflows (*2-in-2-out*) are occluded. Here, flow rate changes above 50% are still observed two generations away from the MSC. In contrast flow rate changes remain below 30% for all capillaries at a MSC with a divergent bifurcation upstream and a convergent bifurcation downstream (*1-in-1-out*). In accordance, the occlusion of a *2-in-2-out* capillary reduces perfusion by nearly 20% in a tissue volume of 200,000 µm^3^. For the occlusion of a capillary with an approximately four times higher baseline flow rate, a 20% drop in perfusion can still be observed for a tissue volume of 500,000 µm^3^. Besides a local decrease in perfusion, single capillary occlusion also causes a decrease in the number of available flow paths in the vicinity of the MSC.

Our observation that the severity of a microstroke is affected by the baseline flow rate of the occluded vessels is in agreement with previous *in vivo* and *in silico* observations for the occlusion of penetrating vessels [2, 4, 16, 28]. Additionally, Nishimura et al. [16] report an RBC velocity reduction in response to single capillary occlusion of 93% and 55% in downstream vessels of generation 1-2 and 3-4, respectively. Although the *in vivo* velocity reductions are slightly higher, they generally compare well with our results for the occlusion of a *2-in-2-out* and a *1-in-2-out* MSC. However, Nishimura et al. [16] do not observe flow reversals and velocity reductions in upstream and parallel vessels. These differences are likely due to the fact that many of the occluded vessels in Nishimura et al. [16] are direct offshoots of the DA main branch. Due to significantly larger flow velocities in the DA main branch, velocity reductions and reversals upstream of the site of occlusion are not to be expected.

To be highlighted is that for all MSC-types the effects of single capillary occlusions are spatially constrained. To be more precise, no significant reduction in flow rate is visible 5 generations away from the MSC and in a tissue volume of 0.3 nl (300,000 µm^3^) around the MSC, the perfusion drops maximally by 10% for all MSC-types. This is in contrast to the occlusion of DAs where the flow rate does not fully recover until the 10^th^ downstream vessel and where the infarct volume is as large as 220 nl [1, 2, 28]. Our results agree qualitatively with previous observations where we studied the impact of single capillary dilations of 10% [40]. As this alteration is significantly smaller than the complete occlusion of a capillary, the flow changes in response to dilation are even limited to capillaries directly adjacent to the dilated vessel.

As the supplied tissue volume of a *1-in-1-out* MSC is 0.056 nl (56,000 µm^3^), which is approximately 15 times smaller than the infarct volume observed for the occlusion of a DA offshoot [2], we postulate that the occlusion of single capillaries does not directly cause tissue damage. This hypothesis is in line with the results of Shih et al. [2]. Nonetheless, our results clearly show that a single capillary occlusion has a strong impact on the local flow field. As such, it seems plausible that the altered flow field is a possible mechanism by which to affect Aβ deposition and clearance [17] or solute clearance via the perivascular space in general [35]. The local disturbances in the flow field and in solute clearance could build-up and cause additional vessel ruptures [6] or occlusions, which subsequently might further impede clearance [37, 57] as well as oxygen and nutrient supply in an increasing area around the MSC. It is important to note that currently the key mechanisms causing the differences in Aβ deposition in response to capillary occlusion are not yet fully understood.

For the occlusion of larger caliber vessels it has been shown that proximal microinfarcts are likely to coalesce [2] and that severe BBB leakage and intravascular platelet aggregation [4] are also observed beyond the microinfarct border. Moreover, Summers et al. [5] report deficits in neuronal activity and functional vasodynamics in response to DA occlusion, which affect an area ∼12-times larger than the microinfarct core. These aspects underline that pathological disturbances are not limited to the core of the microinfarct. However, whether comparable effects can be triggered by the occlusion of a single capillary remains unknown. Likewise, we don’t know whether single capillary occlusion induces local tissue hypoxia or if proximal vessels compensate for the lack of perfusion in the occluded capillary. We suggest that due to the significant effect on the local flow field, single capillary occlusions likely lead to a local drop in tissue oxygenation, which might provoke a cascade of consecutive responses in the affected tissue. The remaining unknowns clearly underline the need for an in-depth *in vivo* quantification of the impact of single capillary occlusion. Based on our results we suggest that the focus of future *in vivo* microstroke studies should be on tissue oxygenation, Aβ deposition and long-term changes in the vicinity of the MSC. In these investigations it is important to keep in mind that the severity of the microstroke is affected by the local vascular topology and the baseline perfusion of the MSC. Consequently, care should be taken that effects are analyzed in an MSC-type specific manner.

As previously stated the severity of a microstroke depends on the local vascular topology. Worthy of note, we observe significant differences in the frequency and the characteristics of the local vascular topologies (MSC-types). *1-in-1-out* is by far the most frequent MSC-type and supplies the largest tissue volume. At the same time it is characterized by having the smallest average flow rate and by containing the smallest number of unique paths connecting DAs and AVs. In contrast *2-in-2-out* is the rarest MSC-type and contains the largest number of flow paths connecting DAs and AVs. We postulate that MSC-type *2-in-2-out* is responsible for distributing blood within the capillary bed and that MSC-type *1-in-1-out* is designed to enable efficient oxygen and nutrient discharge to the tissue.

The frequency of *2-in-2-out* MSCs is low and in a volume of 200,000 µm^3^ around the MSC, 27% of vessels show no flow decrease after a microstroke. These two features suggest that the capillary bed offers an inherent level of robustness towards single capillary occlusion and they agree well with the highly interconnected nature of the capillary bed that allows efficient re-routing of blood flow [26, 28, 41-43, 58, 59].

Nonetheless, the significant differences between the characteristics of the MSC-types raise further questions regarding the origin and the severity of microstrokes. First of all: Would a microstroke be more probable in a *1-in-1-out* MSC? This idea is based on the lower average flow rate in *1-in-1-out* MSCs, which implies that the vessel might be blocked more easily. However, to answer this question we need to improve our understanding of the mechanisms that cause capillary occlusions. It seems likely that occlusions caused by an obstruction are prone to happen in low flow capillaries. However, if plaque deposits or mural cell activity are the origin of capillary occlusions, then the situation is less clear. Secondly: Due to the larger supplied tissue volume, might the occlusion of a *1-in-1-out* MSC be more severe for oxygen and nutrient supply, while the occlusion of a *2-in-2-out* MSC has a larger impact on the flow field? Here, *in vivo* studies monitoring tissue oxygenation in response to capillary occlusion or combined blood flow and oxygen transport simulations could provide insights into the most critical MSC-type for oxygen and nutrient supply.

Furthermore, the impact of a microstroke on tissue oxygenation can be affected by the level of oxygen within the occluded capillary and by the local arrangement of arteriole- and venule-side capillaries with respect to each other. We hypothesized that arteriole-sided capillaries with a high oxygen content might be distributed in a convenient fashion throughout the vasculature to enhance the robustness of oxygen delivery throughout the tissue. Indeed, Nishimura et al. [18] provided supporting evidence for this hypothesis by showing that capillaries with varying topological distances to the DA can be in spatial proximity. However, further studies resolving oxygen partial pressure within capillaries in microvascular networks are necessary to confirm this hypothesis.

Studying the effect of single capillary occlusions in an isolated manner in our *in silico* model is advantageous on the one hand, but limitated on the other hand. For example, our simulation model does not account for dynamic responses of the vasculature. It has been shown that single DA occlusion induces a heterogeneous response in the capillary bed comprising capillary dilations and constrictions [4, 18]. Nonetheless, the *in silico* approach enables us to perform an in-depth study of fluid dynamical changes in response to single capillary occlusion detached from external and internal influences. Moreover, our observations can be conveniently linked to the surrounding vascular topology. These insights can subsequently be used to distinguish changes observed in *in vivo* studies.

Taken together, we show that for 57% of all capillaries an occlusion significantly reduces the flow rate in the adjacent capillaries. Consequently, we conjecture that a single capillary occlusion can be the starting point of a cascade of small consecutive disturbances, which might be relevant for the development of larger microinfarcts and for the progression of pathologies. Importantly, resolving the smallest scale of disturbance is not only essential to improve our understanding of microinfarct development, but might eventually offer novel possibilities for therapeutic treatment and prevention.

## 5 Methods

The presented results are based on a computational model to simulate blood flow in realistic MVNs. The model has been published previously [39] and is briefly revised here. We start by giving a summary of the numerical framework to simulate blood flow in realistic MVNs with tracking of discrete RBCs [39, 48]. Subsequently, we provide more details on the analyses used in the current simulation study.

### 5.1 Blood flow modeling with discrete RBC tracking

The microvascular network is represented as a graph structure, i.e. it consists of a set of nodes *n*_*i*_ connected by a set of edges *e*_*ij*_. The subscript *ij* indicates that edge *e*_*ij*_ is connecting node *n*_*i*_ and *n*_*j*_. Anatomically accurate microvascular networks (MVN) have been acquired by Blinder et al. [28] from the mouse somatosensory cortex by two-photon laser scanning microscopy. They are embedded in a tissue volume of ∼1.6 mm^3^ (MVN1) and ∼2.2 mm^3^ (MVN2) and contain ∼12,100 and ∼19,300 vessels, respectively.

The vessels are labeled as pial arteries (PAs), descending arterioles (DAs), capillaries (Cs), ascending venules (AVs) and pial veins (PVs). For the penetrating vessels, i.e. DAs and AVs, we additionally differ between the main branch and the offshoot vessels. The vessel type is assigned by following the vessels from the cortical surface and by applying a diameter criterion which requests that two subsequent vessels have a diameter smaller than 6 µm in order to change the vessel type from DA to C [39]. The equivalent criterion is applied on the venule side. To differentiate between the main branch of the penetrating vessels and the offshoots we use a criterion that is based on the branching angle and the length of the resulting main branches. This approach ensures that short offshoots are not labeled as main branch.

To compute the pressure field and the blood flow rates in the realistic MVN we employ the continuity equation at every node and Poiseuille’s law along the vessels. This approach is valid due to the small Reynolds numbers in the cortical microvasculature (Re < 1.0 for all vessels). To account for the presence of RBCs the vessel resistance is multiplied by the relative effective viscosity 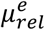. Taken together, Poiseuille’s law reads

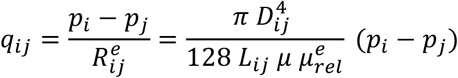

where *D*_*ij*_ and *L*_*ij*_ are the vessel diameter and the length and *P*_*i*_ and *p*_*j*_ are the pressure at node *i*and *j*, respectively. *μ* is the dynamic plasma viscosity and 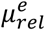 the relative effective viscosity, which is computed as a function of the hematocrit and the vessel diameter as described in Pries et al. (*in vitro* formulation) [60].

The hematocrit of individual vessels is computed from the discretely tracked RBCs. In order to correctly model the motion of RBCs we account for the Fahraeus effect [60, 61] and the phase-separation. The phase-separation in vessels with a diameter larger than 10 µm is described based on the empirical relation by Pries et al. [60]. In vessels with a diameter < 10 µm, single file flow can be assumed and consequently, we postulate that the RBC follows the path of the largest pressure force [39, 48, 62, 63]. The unequal partitioning of RBCs at divergent bifurcations and their impact on the vessel resistance cause a fluctuating flow and pressure field. In the current study we focus on the analysis of the time-averaged flow field of the statistical steady state. Our average is computed over ten turnover times (15.4 s), where one turnover time is defined as the time until 85% of all vessels have been completely perfused at least once.

The pressure boundary conditions are assigned as described in Schmid et al. [39]. In brief, at the pial vessels we make use of existing experimental data and assign a diameter-dependent pressure value. The pressure values at capillary in- and outlets are set based on the simulation results of the hierarchical boundary conditions approach. Here, the realistic MVN is implanted into a large artificial MVN. Subsequently the flow and pressure field for a constant hematocrit is computed and the resulting pressure values are assigned as boundary conditions. The pressure boundary conditions are kept constant for each microstroke scenario.

### 5.2 Microstroke simulations

The microstroke simulations are performed in MVN1. To mimic a microstroke the diameter of a single capillary is set to 0.01 µm. For all investigated scenarios the resulting flow rate in the microstroke capillary (MSC) is < 10^−10^ *μm*^3^*ms*^−1^. The average flow rate in a capillary in MVN1 is 4.*2 μm*^3^*ms*^−1^. This proves that the flow rate in the MSC approaches 0 *μm*^3^*ms*^−1^ and therewith confirms the validity of our microstroke model.

In total there are 11,386 capillaries in realistic MVN1. To ensure that we choose representative MSCs the following selection criteria have to be fulfilled:

1. The MSC should be located in a cylinder with a radius of 444 µm around the x-y-center of the MVN (number of possible MSCs: 8,718).
2. The average flow rate in the MSC has to be > 0.16 *μm*^3^*ms*^−1^ (95% of all capillaries, number of possible MSCs: 8,237). In MVN2 this corresponds to 98% of all capillaries.
3. The average hematocrit needs to be > 0.02 (95% of all capillaries, number of possible MSCs: 7,824).
4. The flow rate in the MSC and its upstream and downstream neighbors should be stable, i.e. frequent flow direction changes should not occur. To be more precise, we allow 5%, 10% and 30% flow direction changes in the MSC, in the first upstream and downstream vessels and in the second and third upstream and downstream vessels, respectively (number of possible MSCs: 6,307). The relative number of flow direction changes is computed by dividing the number of time steps with a flow direction change by the total number of simulated time steps.
5. The MSC is located approximately in the center of the capillary bed, i.e. it is at least three segments apart from the main branch of the DA and AV (number of possible MSCs: 3,565).
6. The MSC has to fit into a bounding box with a volume of 200,000 µm^3^ (number of possible MSC: 3,462).

It should be noted that the “number of possible MSCs” is computed by subsequently considering an additional selection criterion.

One of our goals is to comment on factors influencing the severity of a microstroke. To analyze the impact of different factors, e.g. the baseline flow rate in the MSC, additional selection criteria may be prescribed. These criteria are defined in more detail in the related results sections.

Supplementary Table 1 provides an overview of all selection criteria. In total we analyzed twelve different cases. For each case eight microstroke simulations have been performed.

### 5.3 Thresholded relative change

A major challenge in comparing blood flow simulations in realistic MVNs is the large variety in flow rates, which ranges from 0.09 *μm*^3^*ms*^−1^ to values as large as *26*.*76 μm*^3^*ms*^−1^ in the capillary bed (minimum and maximum of 95% of all flow rates in the capillary bed, median: 1.99 *μm*^3^*ms*^−1^). In such a flow field a large relative change in a vessel with a small baseline flow rate might be negligible, while a small relative change in a vessel with a large baseline flow rate might be significant. To improve the comparability of simulation results, we introduce an absolute threshold, *th*^*abs*^. If the absolute change is smaller than *th*^*abs*^ the relative change is set to 0%.

To choose an appropriate threshold value we compare the average flow rate at two points in time for three different averaging intervals (ten, five and three turnover times). The absolute flow change between the two time points is characteristic for the fluctuations of the baseline flow field. Thus, it can be used as a reference of how large the absolute flow change needs to be such that it is likely to be caused by the microstroke and not by baseline fluctuations. The difference between the two time points is 20 s.

It becomes apparent that for each of the three averaging intervals > 87% of the capillaries change their flow rate by less than 0.1 *μm*^3^*ms*^−1^ (Supplementary Figure 10). Consequently, in the microstroke simulations a flow rate change > 0.1 *μm*^3^*ms*^−1^ is likely caused by the impact of the microstroke and not by baseline fluctuations. Accordingly, we set the absolute threshold *th*^*abs*^ to *μm*^3^*ms*^−1^.

The relative change in flow rate can either be computed directly from the flow rate in the vessel

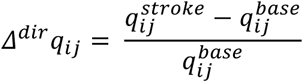

or from the absolute flow rates in the vessel

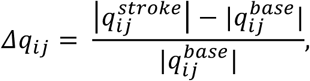

where 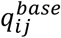and 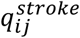 are the flow rates in vessel *ij* for the baseline and the simulation with microstroke, respectively. Even though the second formulation neglects changes in flow direction, it is more suitable for comparing the total perfusion of individual capillaries. As the total perfusion is more relevant for oxygen and nutrient supply, we employ the second expression and analyze flow direction changes separately. Taken together the thresholded relative change is computed as

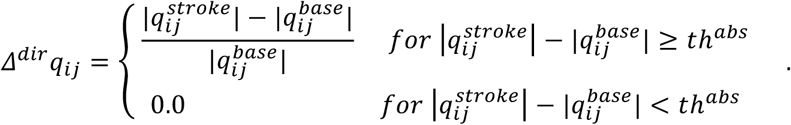

### 5.4 Investigating differences over cortical depth

To analyze differences over cortical depth the realistic MVN is divided into five analysis layers (ALs) each 200 µm thick (Supplementary Figure 4). This analysis approach was first introduced by Schmid et al. [39]. To assign a vessel to an AL, either the source or the target vertex of the vessel has to be within the upper and lower bound of the AL (Supplementary Table 1). The second end point of the vessel is required to be within ±50 µm of the bounds of the AL.

### 5.5 Analysis of total inflow and total flow in an analysis box around MSC

To comment on the blood supply of a tissue volume around the MSC we compute the total inflow into an analysis box around the MSC. The volume of the smallest analysis box is chosen such that each MSC fits into the smallest analysis box. This results in an initial box volume of 200,000 µm^3^, which would be equivalent to a cube with a side length of 58.48 µm. Moreover, for each MSC we have at least 6 capillaries in the initial analysis box and at least 5 capillaries crossing the border of the analysis box. The chosen initial box volume is a compromise between having the smallest possible analysis box around the MSC and ensuring at the same time that sufficient capillaries are within the analysis box to perform a quantitative investigation. The side lengths of the analysis box vary for the different MSC capillaries. To increase the box volume, the side lengths of the smallest analysis box are increased by the same distance in all three dimensions until we reach the desired box volume.

To compute the relative inflow change in response to a microstroke we add up all inflows during baseline and during stroke and calculate the relative difference between the total inflow during baseline and during stroke. It is important to note that due to flow reversals in response to a microstroke the number of inflow vessels can change for the baseline and the microstroke case. The equivalent analysis is repeated for increasing box volumes. The relative inflow change per analysis box is depicted in Figure 2a-d, Supplementary Figure 3c-d and Supplementary Figure 5c-d.

The change in total flow rate per analysis box is computed comparably to the inflow change in an analysis box. Here, instead of computing the total inflow during baseline and during stroke we add up the length weighted total flow rates in the analysis box for baseline and during stroke by summing up the flow rate of all vessels in the analysis box. We consider the vessel tortuosity to compute the vessel length within the analysis box. The total flow rate change per analysis box is used in Figure 2f-h.

### 5.6 Definition of vessels parallel and distant to the MSC

To study the redistribution of flow in an analysis box we introduce three vessel categories: 1) Vessels *upstream and downstream* of the MSC. 2) Vessels that branch off/into an upstream/downstream vessel of generation 1 or 2 of the MSC (*parallel vessels*). Here, we follow each *parallel vessel* of generation 1 three segments downstream/upstream to create the entire set of parallel vessels. 3) All other vessels in the analysis box (i.e. neither upstream, downstream nor parallel vessels) are called *distant vessels*. A schematic drawing of these vessel categories is provided in Figure 2e and Supplementary Figure 2e.

As the whole MVN is connected, *distant vessels* are also connected to the MSC. However, for this vessel category the point of connection is relatively far upstream or downstream. This approach allows us to study changes in response to a microstroke with respect to the topological distance from the MSC. Note that the concept of *parallel vessels* has also been used in Nishimura et al. [16]. However, their definition of *parallel vessels* is different from that used in our analysis. Nishimura et al. [16] consider *parallel vessels* to be only those vessels that have the same source vertex as the occluded vessel.

### 5.7 Computation of the topological supplied tissue volume

To comment on the infarct volume of a microstroke and for further topological studies we compute the supplied tissue volume for each vessel. To do this, the tissue is discretized on a Cartesian grid, in which the realistic MVNs are embedded. One grid cell spans 4 x 4 x 4 µm^3^, which results in ∼11.6 million grid cells for MVN1 and ∼15.3 million grid cells for MVN2. Each cell center is assigned to the closest vessel. By summing over all cells assigned to one vessel we obtain the topological supplied tissue volume per vessel. It is important to note that the topological and the effective supplied tissue volume can differ significantly [51, 52, 64, 65]. This is because of different oxygen levels along the capillary path. Consequently, for vessels with high oxygen levels the effective supplied tissue volume is likely larger than the topological supplied tissue volume and vice versa. Nonetheless, we believe that the topological supplied tissue volume is a representative characteristic for the study of topology and perfusion related aspects of the cortical microvasculature. Please note that for simplicity the topological supplied tissue volume is called supplied tissue volume throughout this manuscript.

### 5.8 Flow paths between descending arterioles and ascending venules

Flow paths between the penetrating vessels are computed by following the flow from the DA to the AV. For this investigation a DA endpoint is defined as the first branch point after the main branch of the DA arteriole. The equivalent definition is used for AVs, i.e. the endpoint of an AV is the point proximal to the capillary bed and the start point of the AV is the root of the penetrating tree at the cortical surface.

To compute all paths between DA and AV we first identify all DA- and AV-endpoints. Subsequently, for each *DA-AV-endpoint-pair* we compute all possible connecting flow paths, which run exclusively through the capillary bed, i.e. if we reach another DA-endpoint before reaching an AV-endpoint, the path is not considered for the *DA-AV-endpoint-pair* under investigation. Note that some DA- and AV-endpoints are not fluid dynamically connected. Additionally, multiple paths enter/leave the MVN across its boundaries. As these paths do not connect a DA with an AV they are not considered any further for this analysis.

The resulting flow path data allows for various investigations:

1. The computation of the total number of flow paths in the MVN (Supplementary Figure 9b).
2. The computation of the number of flow paths per capillary and how this number changes during a microstroke (Figure 4c, Supplementary Figure 9c). In Figure 4c we are interested in the number of paths that persist in the MSC during the microstroke. The relative number of paths is computed as 100%. 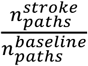 In Supplementary Figure 9c we focus on the relative change in the number of paths during baseline and during stroke. This is calculated as 100%. 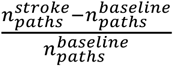, where 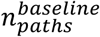 and 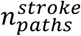are the numberof flow paths through an individual capillary during baseline and during stroke, respectively.
3. Instead of looking directly at the flow paths, we can also analyze the number of unique *DA-AV-endpoint-pairs*. Here, we can either look at the total number of unique *DA-AV-endpoint pairs* (Supplementary Figure 9d) or we can count the number of unique *DA-AV-endpoint-pairs* that are connected by a path through a predefined capillary (Supplementary Figure 9e). As before, this quantity can be compared between the baseline and the stroke simulation. In Supplementary Figure 9d we compare the difference of the total number of *DA-AV-endpoint pairs* before and after stroke. In Supplementary Figure 9e we show the absolute number of *DA-AV-endpoint-pairs* connected by a path through the MSC before and after stroke.
4. Lastly, we count the number of unique flow paths between a given *DA-AV-endpoint-pair* (Figure 4e). To comment on the redistribution of flow with respect to the MSC we introduce three categories to classify *DA-AV-endpoint-pairs* (Figure 4d): Category 1) before and after stroke there is at least one path that leads from the DA- to the AV-endpoint through the MSC; Category 2) only before stroke is there at least one path that leads from the DA- to the AV-endpoint through the MSC and Category 3) none of the paths between the given *DA-AV-endpoint-pair* go through the MSC. Each *DA-AV-endpoint-pair* is assigned to the according category and the ratio of the number of unique flow paths is computed 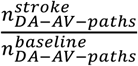, where 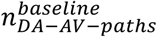and 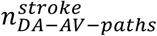are the number of unique flow paths between a given *DA-AV-endpoint-pair* during baseline and during stroke.

### 5.9 Computation of *AV-factor*

The *AV-factor* is computed to distinguish between capillaries that are close to DAs (arteriole-sided capillaries) and those that are close to AVs (venule-sided capillaries). For each capillary *ij* we computed all paths to all DA endpoints and all paths to all AV endpoints. This results in a set of path lengths on the arteriole side of the capillary 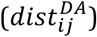 and a set of path lengths on the venule side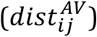. To compute the *AV-factor* for capillary *ij* we use the median of each set of path lengths, i.e.

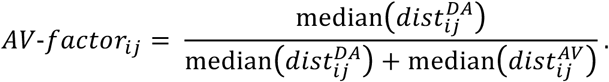

The resulting *AV-factor* lies between 0 and 1 and approaches 0 on the arterial side and 1 on the venule side of the capillary path.

## 6 Acknowledgements

We thank David Kleinfeld, Philbert Tsai and Pablo Blinder for sharing the realistic microvascular networks with us. Moreover, we are grateful for the fruitful discussions of our results with Eva Erlebach, Robert Epp and Jacqueline Condrau. Additionally, we thank Eva Erlebach for her feedback on our manuscript. We thank Karen Everett for editorial help with the manuscript.

## 7 Competing interests

The authors declare that no competing interests exist.

## 9 Supplementary information

**Supplementary Figure 1.**
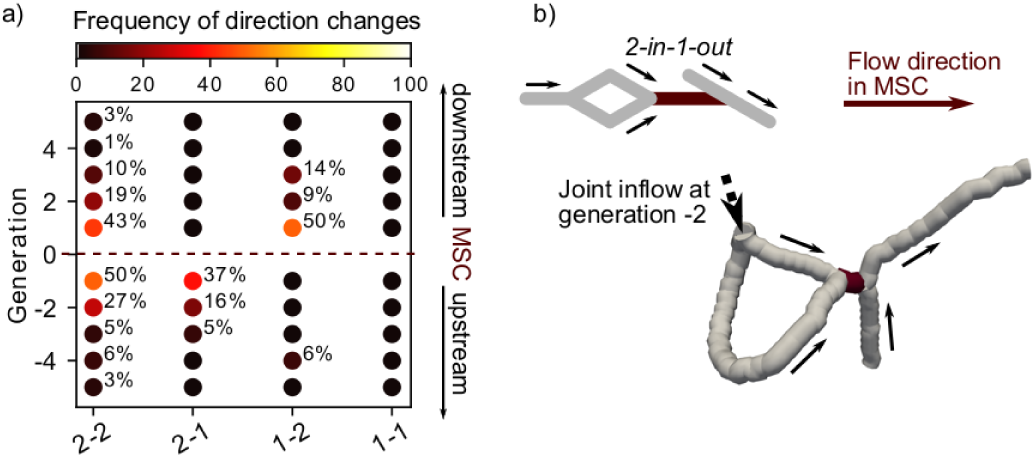
Flow direction changes for the four microstroke capillary (MSC) types. a) Percentage of flow direction changes for the four possible topological configurations at a MSC (Figure 1a-d). Only percentage values >0% are annotated. For each topological configuration eight microstroke simulations have been performed. The percentage is computed by identifying all vessels with a flow direction change for each generation across all eight microstroke simulations and setting it into relation to the total number of vessel per generation. Abbreviations of the four MSC-types: 2-2: *2-in-2-out*, 2-1: *2-in-1-out*, 1-2: *1-in-2-out*, 1-1: *1-in-1-out*. b) Illustration of the specific vascular configuration for the *2-in-1-out* MSC that leads to a cessation of flow in the generation −1 vessels. A schematic (upper left) and a realistic example (lower right) are provided. The MSC (red) and its adjacent vessels (grey, generation −1 and 1) are depicted. The schematic also shows the joint inflow at generation −2.

**Supplementary Figure 2.**
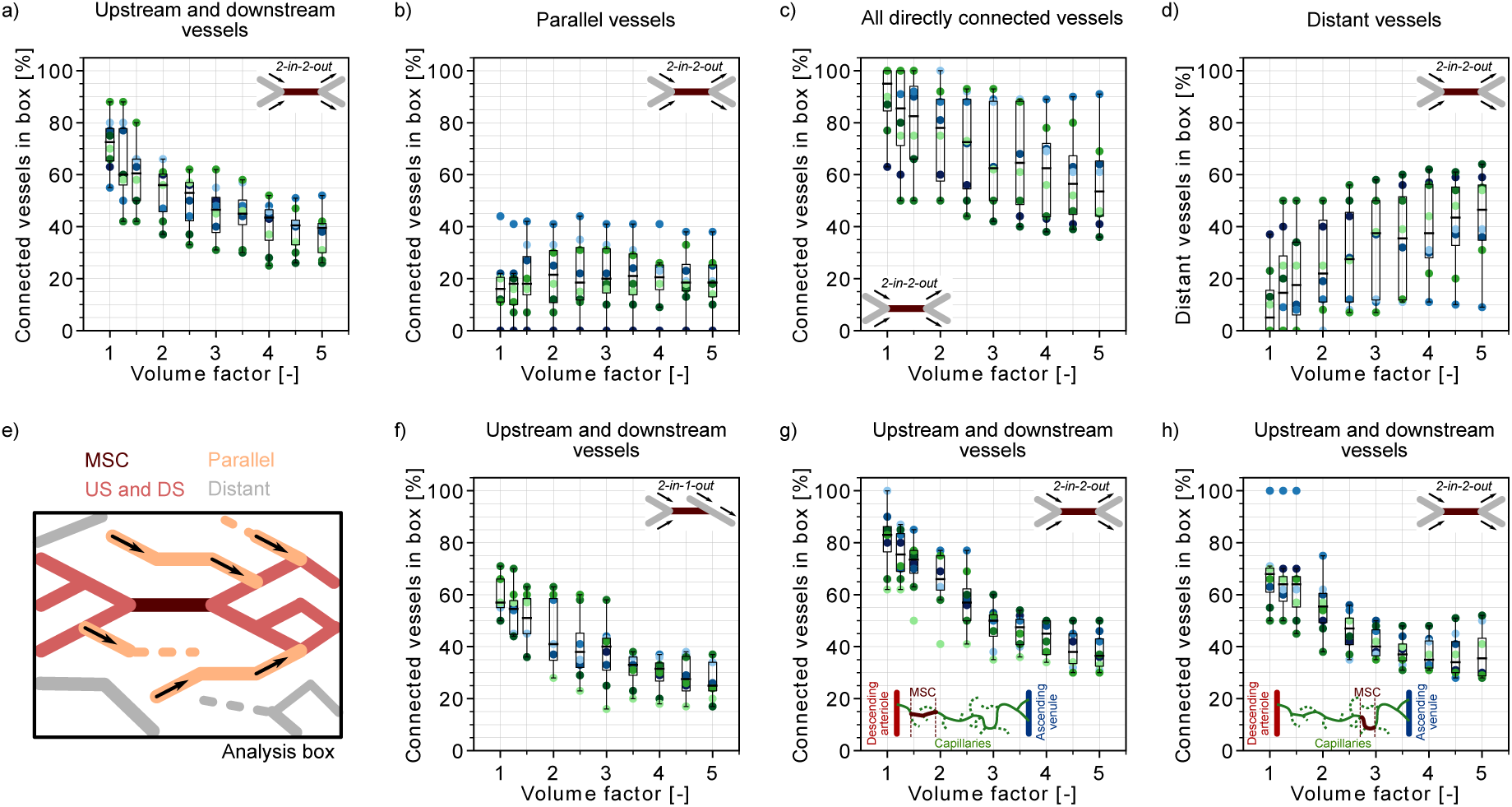
Occurrences of different vessels categories within the analysis box around the microstroke capillary (MSC). a)-d) Percentage of vessel within the analysis box which are positioned differently with respect to the *2-in-2-out* MSC. a) Percentage of vessels upstream and downstream of the MSC for an increasing analysis box volume. b) Percentage of parallel vessels (Methods) c) Percentage of vessels directly connected to the MSC, i.e. upstream, downstream and parallel vessels. d) Percentage of distant vessels, i.e. vessels that are neither upstream, downstream nor parallel. e) Schematic to illustrate the concept of vessels upstream, downstream, parallel and distant to the MSC (Methods). f)-h) Percentage of upstream and downstream vessels of the MSC for an increasing analysis box volume for different cases. f) *2-in-1-out*, g) *2-in-2-out* close to a descending arteriole (DA) and h) *2-in-2-out* far away from a DA. Further details on the selection criteria are provided in Supplementary. The initial box volume, i.e. volume factor = 1, is 200,000 µm^3^. It is chosen such that each MSC fits into the initial analysis box and that each analysis box has at least five inflows. The eight microstrokes per case are depicted by the color-coded spheres. The boxplots are based on the available data for each generation.

**Supplementary Figure 3.**
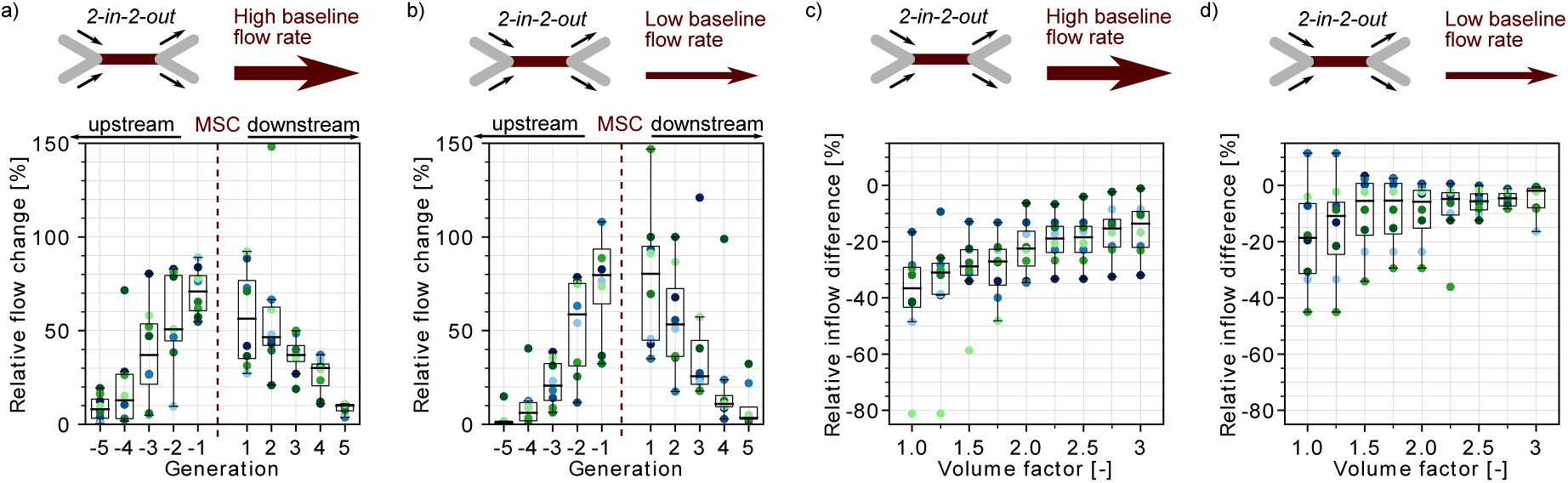
Impact of the baseline flow rate on the severity of a microstroke in a *2-in-2-out* microstroke capillary (MSC). a)-b) Average relative change in flow rate ***Δq***_***ij***_ for capillaries up- and downstream of the MSC. c)-d) Relative inflow difference for an increasing box volume around the MSC. The initial box volume, i.e. volume factor = 1, is 200,000 µm^3^. It is chosen such that each MSC fits into the initial analysis box and that each analysis box has at 5 inflow vessels. The relative inflow difference is computed by summing up the inflows across the borders of the box for the baseline and the stroke simulation (Methods). a) and c) show the results for a high baseline flow rate (7.0-25.0 µm^3^ms^−1^) and b) and d) for a lower baseline flow rate (0.1-4.0 µm^3^ms^−1^). Further details on the selection criteria are provided in Supplementary. b) and d) are also depicted in Figure 1a and 2a and are only provided to facilitate comparison between the two cases. The eight microstrokes per case are depicted by the color-coded spheres. The boxplots are based on the available data for each generation.

**Supplementary Figure 4.**
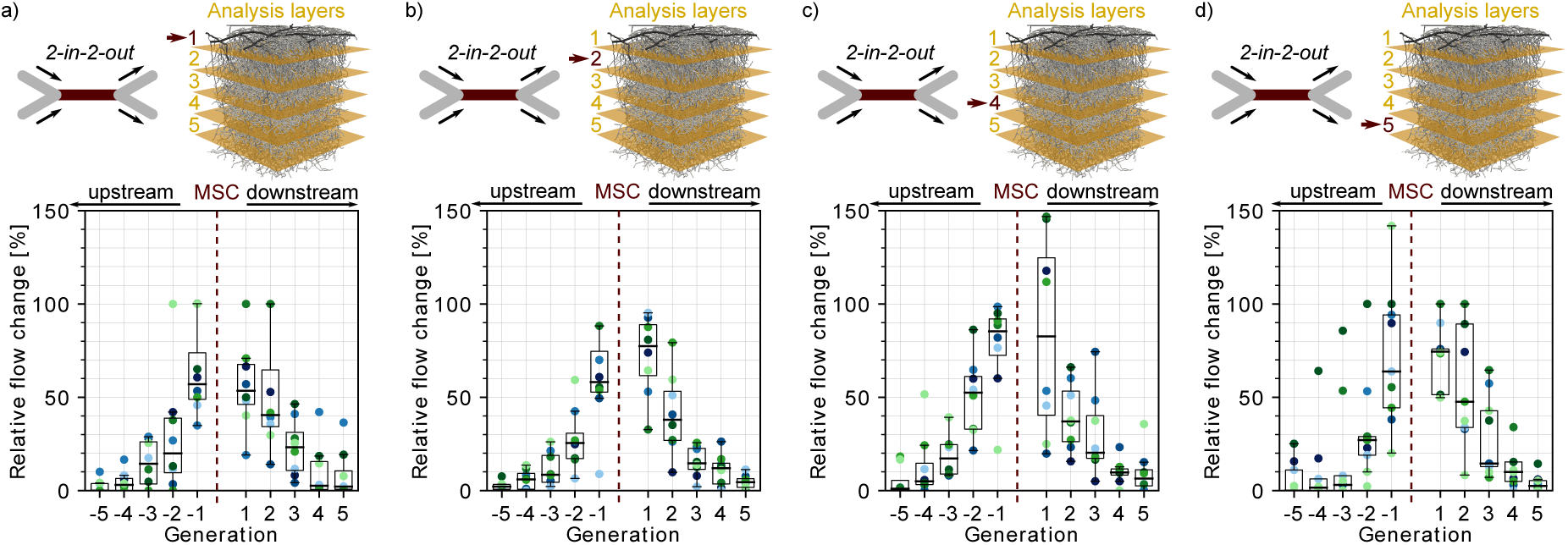
Impact of the cortical depth on the severity of a microstroke in a *2-in-2-out* microstroke capillary (MSC). a)-d) Upper panel: Schematic of a *2-in-2-out* and the realistic microvascular network, which has been divided into 5 analysis layers (AL) each 200 µm thick. The arrow indicates the AL for which the results are depicted below. Lower panel: Average relative change in flow rate ***Δq***_***ij***_ for capillaries up- and downstream of the MSC. For each cortical depth the flow field for eight MSCs has been computed. The average relative change per generation for each of the eight simulations is depicted by the **color-coded** spheres. The boxplots are based on all available data for each generation. Further details on the selection criteria are provided in Supplementary Table 1.

**Supplementary Figure 5.**
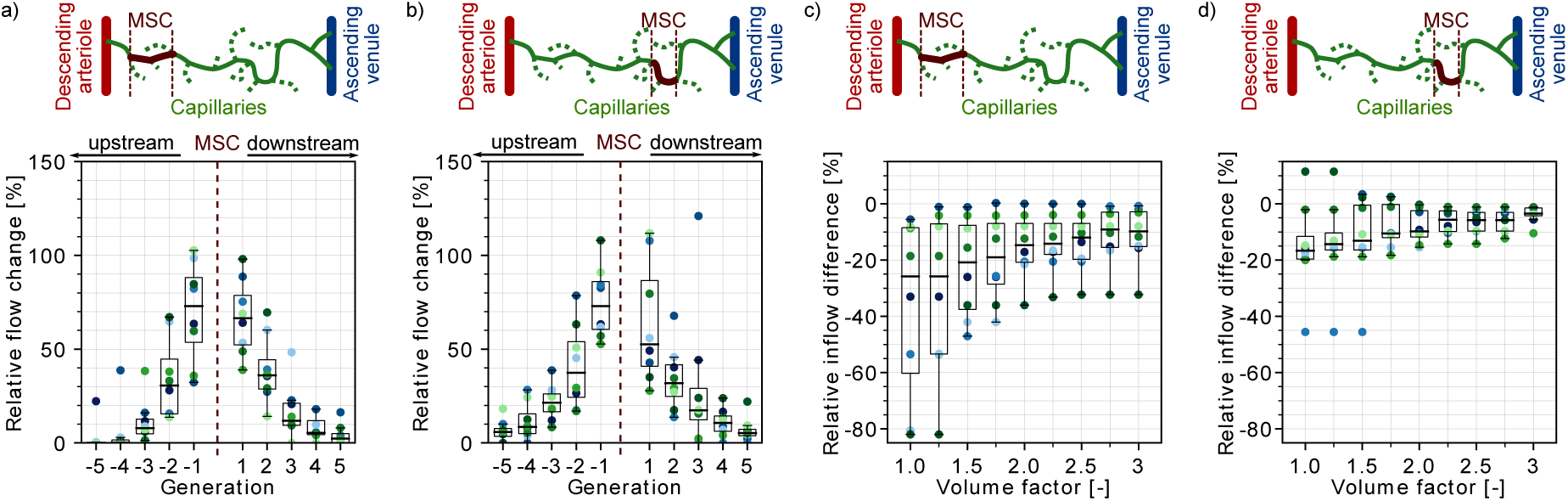
Impact of the distance of the microstroke capillary (MSC) to the penetrating vessels on the severity of a microstroke in a *2-in-2-out*. a)-b) Average relative change in flow rate ***Δq***_***ij***_ for capillaries up- and downstream of the MSC. c)-d) Relative inflow difference for an increasing box volume around the MSC. The upper panel shows a schematic of the location of the MSC capillary along an exemplary capillary path between descending arteriole (DA) and ascending venule (AV). The initial box volume, i.e. volume factor = 1, is 200,000 µm^3^. It is chosen such that each MSC fits into the initial analysis box and that each analysis box has five inflows. The relative inflow difference is computed by summing up the inflows across the borders of the analysis box for the baseline and the stroke simulation (Methods). a) and c) show the results for MSC close to the DA and b) and d) for MSCs distant to the DAs. It should be noted that the larger inflow rate reduction for vessels close to DAs (c)) is not caused by an overall larger reduction in flow rate but by a high percentage of upstream and downstream vessel in the analysis box (Supplementary Figure 2g-h). As described in the main text the flow decrease is largest in upstream and downstream vessels and consequently a large number of upstream and downstream vessels in the analysis box will reflect in a larger inflow reduction. Further details on the selection criteria are provided in Supplementary. The eight microstrokes per case are depicted by the color-coded spheres. The boxplots are based on the available data for each generation.

**Supplementary Figure 6.**
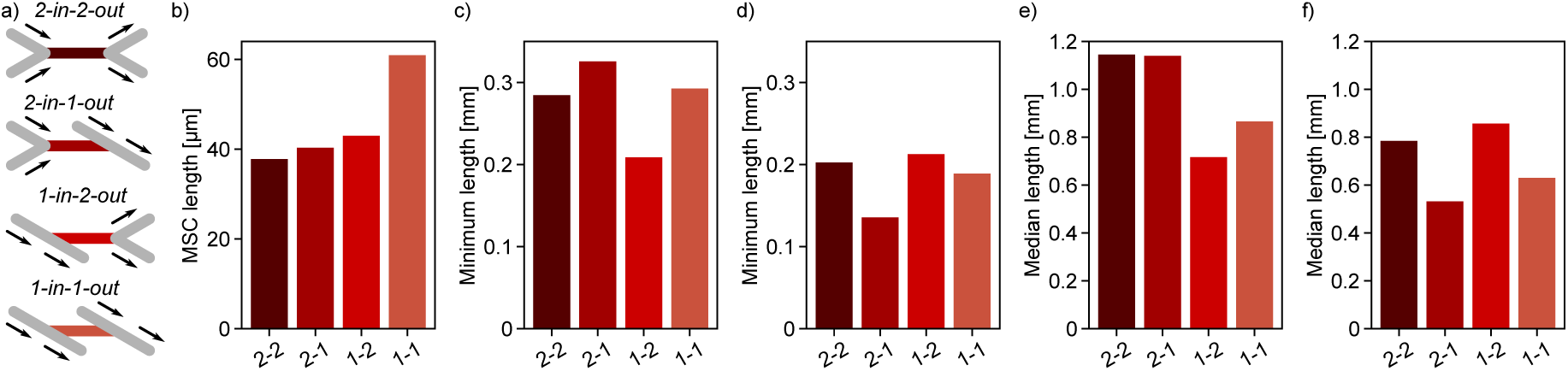
Length and distance to penetrating vessels of microstroke capillaries (MSC) of different types. a) Schematic of the four topological configurations at the MSC. The MSC is color coded in accordance with subfigures b)-f). b) Median vessel length of the four MSC types. c)-d) Median length of the minimum distance of all paths leading from a MSC to descending arteriole (DA, c)) and ascending venule (AV, d)) main branches. e)-f) Median length of the median distance of all paths leading from a MSC to DA (e) and AV (f) main branches. Abbreviations of the four MSC-types: 2-2: *2-in-2-out*, 2-1: *2-in-1-out*, 1-2: *1-in-2-out*, 1-1: *1-in-1-out*. The statistics are based on all capillaries that fulfill the general selection criteria described in the Methods. The fifth selection criterion is less strict for the current analysis, i.e. the capillary only has to be one segment apart from the DA/AV, and the sixth criterion is not applied. This results in 4818 and 8544 capillaries for analysis for MVN1 and MVN2, respectively. First the median within each MVN is computed and subsequently the average across the two MVNs.

**Supplementary Figure 7.**
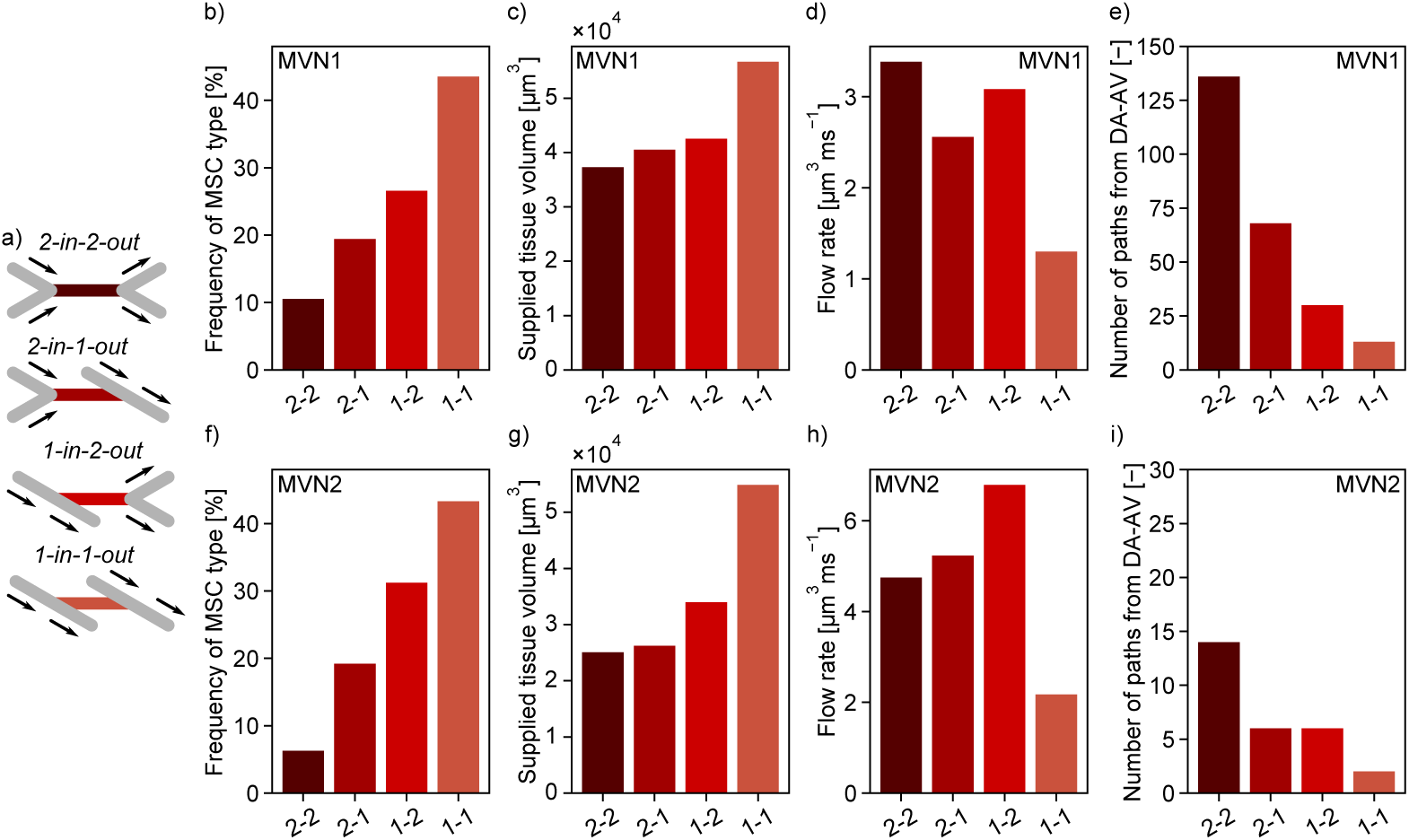
Comparison of the characteristics of the four topological configurations at a microstroke capillary (MSC) for MVN1 and MVN2. a) Schematic of the four topological configurations at the MSC. The MSC is color coded in accordance with subfigures b)-i). b), f) Frequency of occurrence of the four MSC types for MVN1 and MVN2, respectively. c), g) Median supplied tissue volume for the four MSC types for MVN1 and MVN2 (Methods). d), h) Median flow rate the four MSC types for MVN1 and MVN2, respectively. e), i) Median number of unique paths leading through a MSC from the descending arteriole (DA) to the ascending venule (AV) for MVN1 and MVN2, respectively. Abbreviations of the four MSC-types: 2-2: *2-in-2-out*, 2-1: *2-in-1-out*, 1-2: *1-in-2-out*, 1-1: *1-in-1-out*. The statistics are based on all capillaries that fulfill the general selection criteria described in the Methods. The fifth selection criterion is less strict for the current analysis, i.e. the capillary only has to be one segment apart from the DA/AV, and the sixth criterion is not applied. This results into 4818 and 8544 capillaries for analysis for MVN1 and MVN2, respectively.

**Supplementary Figure 8.**
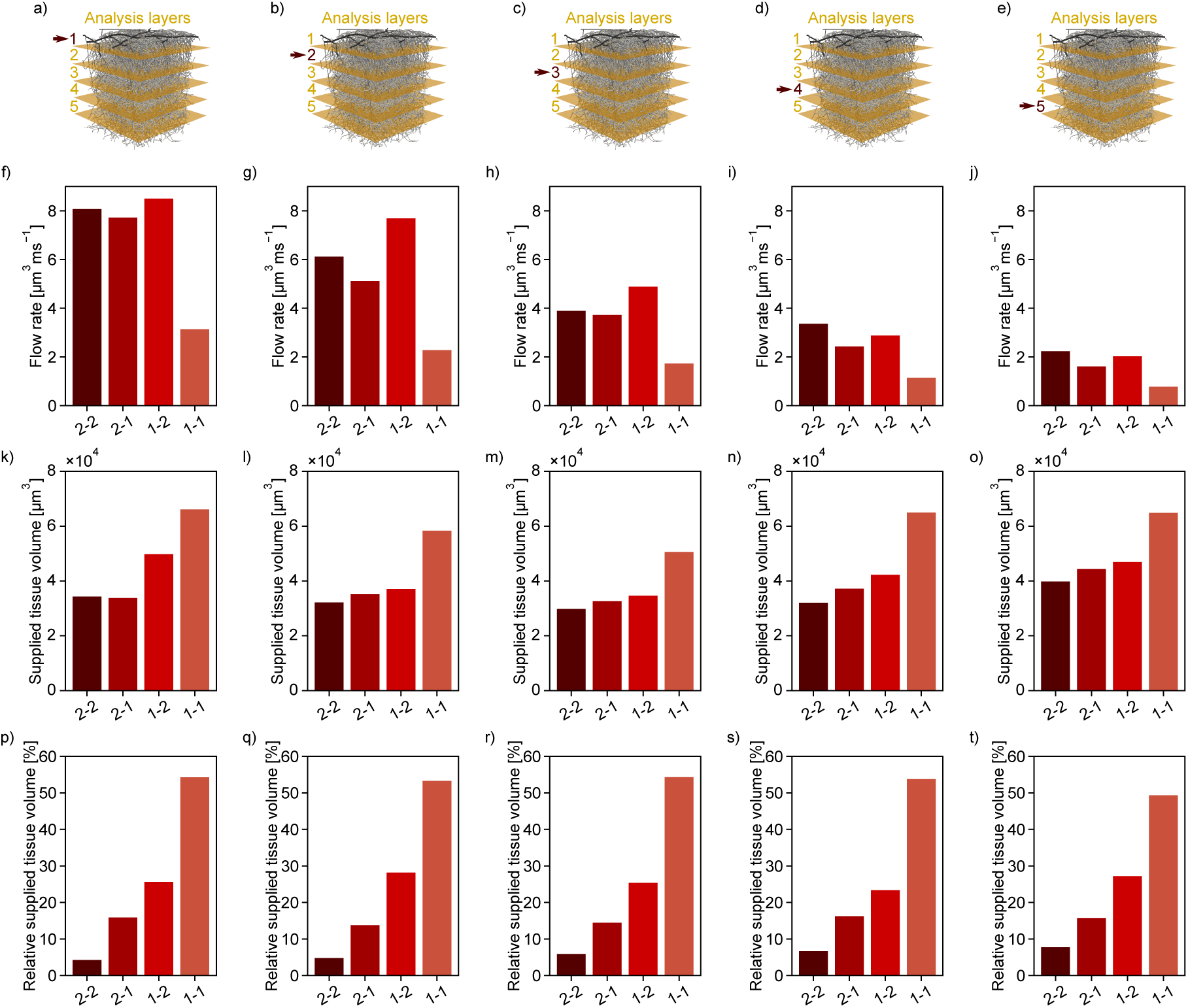
Median flow rate, median and total relative supplied tissue volume for the four microstroke capillary (MSC) types over cortical depth. a)-e) Schematic of a realistic microvascular network, which has been divided into 5 analysis layers (AL) each 200 µm thick. The arrow indicates for which AL the results are depicted in the subplots below. f)-j) Median flow rate for the different MSC-types over cortical depth. k)-o) Median supplied tissue volume for the different MSC-types over cortical depth (Methods). p)-t) Total relative supplied tissue volume for the different MSC-types over cortical depth. The relative supplied tissue volume is calculated by summing up the supplied tissue volume for each MSC-type and dividing it by the total tissue volume per AL. Abbreviations of the four MSC-types: 2-2: *2-in-2-out*, 2-1: *2-in-1-out*, 1-2: *1-in-2-out*, 1-1: *1-in-1-out*. The statistics are based on all capillaries that fulfill the general selection criteria described in the Methods. The fifth selection criterion is less strict for the current analysis, i.e. the capillary only has to be one segment apart from the DA/AV, and the sixth criterion is not applied. First the median within each MVN is computed and subsequently the average across the two MVNs.

**Supplementary Figure 9.**
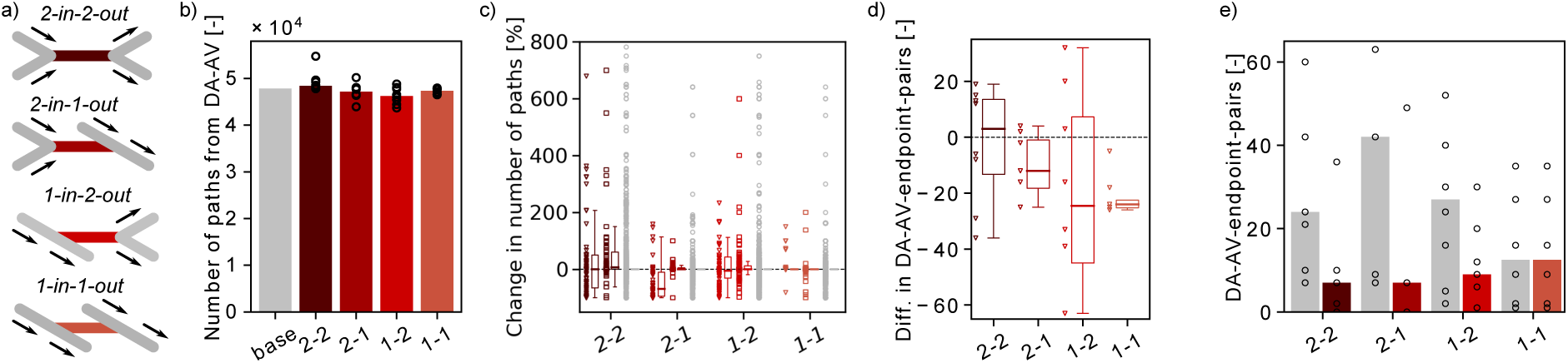
Changes in the number of flow paths and the number of *DA-AV-endpoint-pairs* in response to a microstroke. a) Schematic of the four topological configurations at the microstroke capillary (MSC). The MSC is color coded in accordance with subfigures b)-e). b) Total number of flow paths connecting a descending arteriole (DA) to an ascending venule (AV) in MVN1. The bar plot depicts the results for the baseline simulation (base) and the median for the each microstroke case. The spheres show the total number of flow paths for the eight microstroke simulations per MSC-type. c) Relative change in the number of flow paths through upstream and downstream, parallel and distant of capillaries (Methods). The relative change is computed from the total number of paths during baseline and during stroke (Methods). Only capillaries that are at least along one flow path between DA and AV during baseline are considered. The data of all eight microstroke cases per MSC-type is shown in the scatter points and summarized in the boxplot right of it. Triangles: Capillaries upstream and downstream of the MSC (3 generations). Squares: Capillaries parallel to the MSC. Circles: Capillaries distant to the MSC. 0.1% of the data is not displayed (change > 800%). d) Difference (Diff.) in the total number of unique *DA-AV-endpoint pairs* (Methods, Diff. < 0: Decrease in the number of unique *DA-AV-endpoint pairs* with respect to baseline). The data of all eight microstroke cases per MSC-type is shown in the scatter points and summarized in the boxplot right of it. For MSC-types *2-in-1-out, 1-in-2-out* and *1-in-1-out* only 7 microstroke cases are shown in the scatter points because each of these types has a microstroke case with a difference < −90. e) Number of *DA-AV-endpoint-pairs*, which are connected by at least on path through the MSC (category 1). The bar plot depicts the median for the each MSC type. The grey bar shows the median during baseline and the red bars the median during stroke. The spheres show the number of *DA-AV-endpoint-pairs* for the eight microstrokes for each MSC-type. Note that three data points have more than 90 *DA-AV-endpoint-pairs* and are not displayed by the spheres. Abbreviations of the four MSC-types: 2-2: *2-in-2-out*, 2-1: *2-in-1-out*, 1-2: *1-in-2-out*, 1-1: *1-in-1-out*.

**Supplementary Figure 10.**
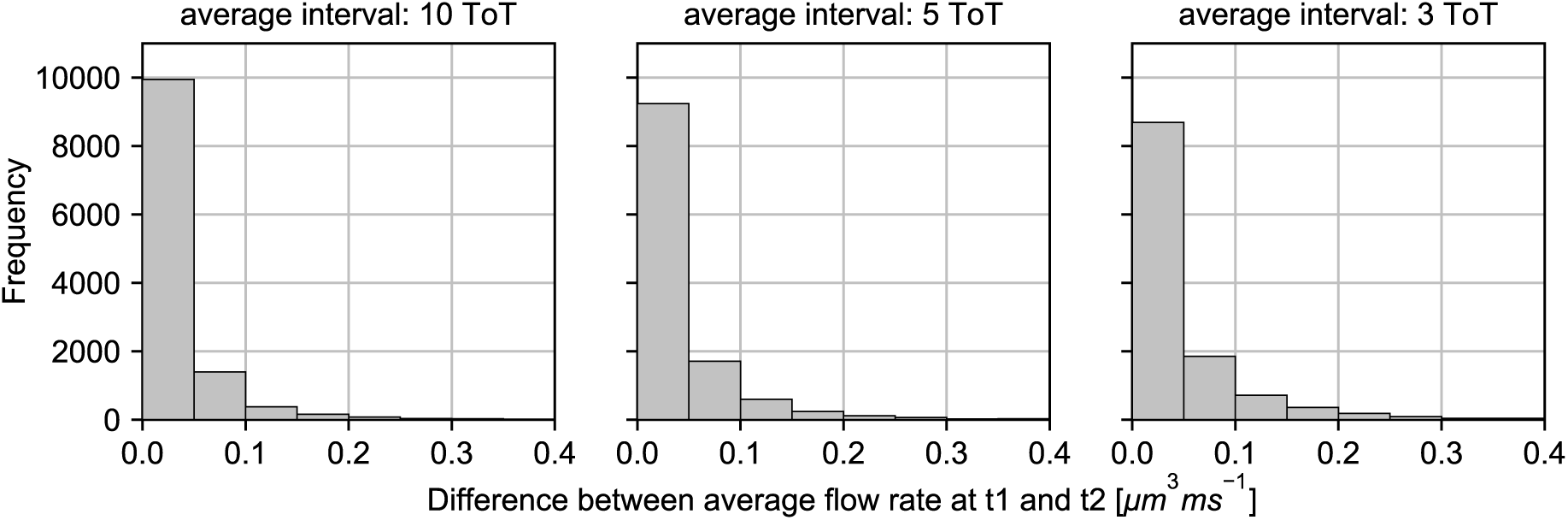
Absolute differences between averaged flow rates in all capillaries at two time points t1 and t2. The time difference between the two time points is 20s. In the left panel the absolute differences for an averaging interval of 10 turnover times (ToT) are displayed. In the middle and the left panel the differences for averaging intervals of 5 ToTs and 3 ToTs are shown. The absolute differences between the averaged results increase for smaller averaging intervals. For an averaging interval of 10 ToT for 94% of all vessels the absolute difference is smaller than 0.1 µm^3^ms^−1^. This value decreases to 91% and 87% for an averaging interval of 5 ToT and 3 ToT, respectively.

**Supplementary Table 1.** Overview of the eight selection criteria used to analyze the impact of structural and functional characteristics on the severity of a microstroke. The different microstroke capillary (MSC) types are depicted in Figure 1a-d. For cases 1-7 the cortical depth selection criterion requires that only the source of the MSC be within the given range. For cases 8-12 at least one of the vertices should be within the given range, while the second one may be ±50 µm outside the given range. The mean and standard deviation (std) are calculated from the results of the baseline simulation for the eight chosen MSC per case. For the mean and std of the cortical depth the values of the source and the target vertex are both considered. The definition of the main branch is provided in the methods. DA: descending arteriole. AV: ascending venule.

| Case | MSC type | Flow rate [$\mu\text{m}^3\text{ms}^{-1}$] | | | Cortical depth [$\mu\text{m}$] | | | Vessels to main branch of DA/AV |
| --- | --- | --- | --- | --- | --- | --- | --- | --- |
| | | min | max | mean ± std | min | max | mean ± std | |
| 1 | 2-in-2-out | 0.1 | 4.0 | $2.87 \pm 0.70$ | 400 | 700 | $557 \pm 70$ | > 3 (DA), >3 (AV) |
| 2 / 3 | 2-in-1-out | 0.1 | 4.0 | $2.32 \pm 0.81$ | 400 | 700 | $597 \pm 111$ | > 3 (DA), >3 (AV) |
| 3 / 4 | 1-in-2-out | 0.1 | 4.0 | $3.03 \pm 1.12$ | 400 | 700 | $520 \pm 70$ | > 3 (DA), >3 (AV) |
| 4 / 6 | 1-in-1-out | 0.1 | 4.0 | $1.35 \pm 0.41$ | 400 | 700 | $555 \pm 108$ | > 3 (DA), >3 (AV) |
| 5 / 2 | 2-in-2-out | 7.0 | 25.0 | $10.7 \pm 3.55$ | 400 | 700 | $518 \pm 64$ | > 3 (DA), >3 (AV) |
| 6 / 5 | 2-in-2-out | 0.1 | 4.0 | $2.89 \pm 0.56$ | 400 | 700 | $611 \pm 82$ | 2-3 (DA), >3 (AV) |
| 7 | 2-in-2-out | 0.1 | 4.0 | $2.67 \pm 0.90$ | 400 | 700 | $542 \pm 110$ | > 7 (DA), >3 (AV) |
| 8 | 2-in-2-out | 0.2 | 4.2 | $2.41 \pm 1.23$ | 0 | 200 | $122 \pm 64$ | > 3 (DA), >3 (AV) |
| 9 | 2-in-2-out | 0.2 | 4.2 | $2.81 \pm 0.89$ | 200 | 400 | $350 \pm 48$ | > 3 (DA), >3 (AV) |
| 10 | 2-in-2-out | 0.2 | 4.2 | $2.87 \pm 0.90$ | 400 | 600 | $506 \pm 50$ | > 3 (DA), >3 (AV) |
| 11 | 2-in-2-out | 0.2 | 4.2 | $2.63 \pm 0.86$ | 600 | 800 | $688 \pm 43$ | > 3 (DA), >3 (AV) |
| 12 | 2-in-2-out | 0.2 | 4.2 | $1.54 \pm 0.87$ | 800 | 100 | $894 \pm 50$ | > 3 (DA), >3 (AV) |

**Supplementary Table 2.** Distribution of microstroke capillary (MSC) types over cortical depth for microvascular network (MVN) 1 and 2. AL: Analysis Layer. Abbreviations of the four MSC-types: 2-2: *2-in-2-out*, 2-1: *2-in-1-out*, 1-2: *1-in-2-out*, 1-1: *1-in-1-out*.

| MVN | AL | Cortical depth [$\mu\text{m}$] | | Total number of MSC | Frequency of different MSC types [%] | | | |
| --- | --- | --- | --- | --- | --- | --- | --- | --- |
|  |  | min | max |  | 2-2 | 2-1 | 1-2 | 1-1 |
| 1 | 1 | 0 | 200 | 844 | 8.1 | 21.7 | 23.7 | 46.5 |
| 1 | 2 | 200 | 400 | 1005 | 8.2 | 15.6 | 31.7 | 44.5 |
| 1 | 3 | 400 | 600 | 1137 | 10.4 | 18.8 | 26.0 | 44.8 |
| 1 | 4 | 600 | 800 | 600 | 11.9 | 21.0 | 23.7 | 43.4 |
| 1 | 5 | 800 | 1000 | 805 | 12.2 | 20.1 | 27.5 | 40.3 |
| 2 | 1 | 0 | 200 | 1549 | 5.0 | 20.2 | 31.1 | 43.6 |
| 2 | 2 | 200 | 400 | 1916 | 4.4 | 17.9 | 35.2 | 42.4 |
| 2 | 3 | 400 | 600 | 2047 | 6.2 | 18.6 | 31.5 | 43.7 |
| 2 | 4 | 600 | 800 | 1819 | 7.3 | 19.8 | 28.6 | 44.3 |
| 2 | 5 | 800 | 1000 | 1318 | 8.8 | 17.4 | 30.0 | 43.8 |

**Supplementary Table 3.**
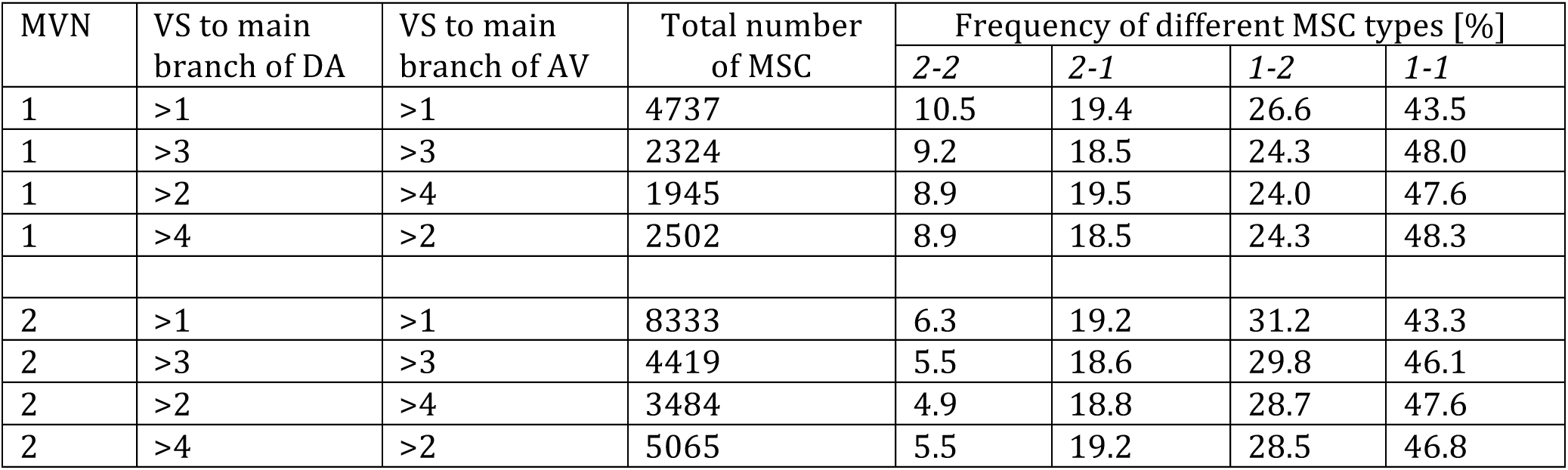
Distribution of microstroke capillary (MSC) types along the path between descending arteriole (DA) and ascending venule (AV) for microvascular network (MVN) 1 and 2. VS: Vessel segments. Abbreviations of the four MSC-types: 2-2: *2-in-2-out*, 2-1: *2-in-1-out*, 1-2: *1-in-2-out*, 1-1: *1-in-1-out*.

